# Design, synthesis and cellular characterization of a new class of IPMK kinase inhibitors

**DOI:** 10.1101/2024.05.09.593371

**Authors:** Yubai Zhou, Pratima Chapagain, Desmarini Desmarini, Dilipkumar Uredi, Lucia E. Rameh, Julianne T. Djordjevic, Raymond D. Blind, Xiaodong Wang

## Abstract

Many genetic studies have established the kinase activity of inositol phosphate multikinase (IPMK) is required for the synthesis of higher-order inositol phosphate signaling molecules, the regulation of gene expression and control of the cell cycle. These genetic studies await orthogonal validation by specific IPMK inhibitors, but no such inhibitors have been synthesized. Here, we report complete chemical synthesis, cellular characterization, structure-activity relationships and rodent pharmacokinetics of a novel series of highly potent IPMK inhibitors. The first-generation compound **1** (UNC7437) decreased cellular proliferation and tritiated inositol phosphate levels in metabolically labeled human U251-MG glioblastoma cells. Compound **1** also regulated the transcriptome of these cells, selectively regulating genes that are enriched in cancer, inflammatory and viral infection pathways. Further optimization of compound 1 eventually led to compound **15** (UNC9750), which showed improved potency and pharmacokinetics in rodents. Compound **15** specifically inhibited cellular accumulation of InsP_5_, a direct product of IPMK kinase activity, while having no effect on InsP_6_ levels, revealing a novel metabolic signature detected for the first time by rapid chemical attenuation of cellular IPMK activity. These studies designed, optimized and synthesized a new series of IPMK inhibitors, which reduces glioblastoma cell growth, induces a novel InsP_5_ metabolic signature, and reveals novel aspects inositol phosphate cellular metabolism and signaling.

## INTRODUCTION

The human inositol phosphate multikinase (IPMK) is a promiscuous nuclear kinase that can phosphorylate several species of inositol phosphates (1–4) as well as the membrane phospholipid PI(4,5)P2 (5, 6), which are all biologically important metabolites and/or small signaling molecules. IPMK generates several InsP_5_ species, several InsP_4_ species, is genetically required for full InsP_6_ production (1, 2), generates PIP_3_ in membrane systems (2, 5) and can phosphorylate PI(4,5)P_2_ bound to a transcription factor (6). As might be expected with such a diverse set of activities, genetic loss of IPMK results in large changes to many important cellular processes, including cell proliferation and gene expression. Genetic studies suggest the kinase activity of IPMK controls many aspects of these complex processes, but orthogonal chemical validation of the genetic experiments had awaited development of specific inhibitors of IPMK. The lack of IPMK inhibitors prevents orthogonal chemical validation of the promiscuity of IPMK kinase activity, further investigation into the cellular kinetics with which these metabolites are synthesized, and the functional consequences of their synthesis in cells.

Genetic loss of IPMK is embryonic lethal in flies and mice (7, 8), suggesting clinical use of IPMK inhibitors might carry detrimental side effects. However, several lines of evidence suggest IPMK inhibitors will be well tolerated in humans. Flies and mice heterozygous for IPMK loss bear no detectable phenotypes (7, 8) and mutant hypomorphic IPMK alleles from plants can rescue developmental loss of IPMK in animals (7, 9). Orthologous IPMK alleles from various species can rescue loss of IPMK function in animal development, even when the plant *Arabidopsis thaliana* IPMK is used to rescue (7). These data all suggest that only a small fraction of the total IPMK activity is required during embryonic development of animals, and further that the protein sequence providing the activity matters little, if at all, during embryonic development of animals. Tissue-specific developmental knockouts of IPMK driven by CRE-promoters in mice results in viable offspring in all cases, liver, adipocytes and brain have tissue-specific IPMK knockout mice that have revealed important physiology (10–12), but show no increased mortality and relatively minor phenotypes. The developmental defect in IPMK-knockout mice occurs early in development (E9.5) specifically during neural tube closure (8). Thus, the genetic data suggest that some small amount of IPMK activity is absolutely required during vertebrate development, but the data also suggest IPMK loss should be well tolerated in adult animals. Of course, that conclusion awaits chemical validation using IPMK inhibitors in pre-clinical adult animal models.

IPMK has also been linked to cancer through several studies (5, 13–20). Loss of IPMK kinase activity in cells is associated with decreased cellular proliferation, thus inhibitors against IPMK have been speculated to have utility as potential anti-cancer therapies. There is also evidence suggesting that inhibiting IPMK could be therapeutic in particular phosphatase and tensin homolog (PTEN)-negative cancers that are independent of AKT (6). IPMK localizes to the nucleus where it can activate gene expression in a kinase-dependent manner (5, 6, 21–23). In some instances, IPMK gene-expression activity has been shown to be antagonized by the tumor suppressor PTEN in a phosphatase-dependent manner, with IPMK functioning as the “on” switch and PTEN the opposing “off” switch in these transcriptional studies (6).

Transcriptional activation by IPMK was not recapitulated by the classic PI3K p110α, and wortmannin did not recapitulate the effects of attenuating IPMK activity, together suggesting PTEN is not antagonizing p110 PI3-Kinase activity, but rather PTEN was antagonizing the activity of IPMK (6). This phosphatase-dependent nuclear PTEN activity has clear clinical implications - if the growth of PTEN-negative tumors is being driven by the function of nuclear PTEN in transcriptional regulation (24), those tumors might be expected to respond to an IPMK inhibitor, but not respond to inhibitors of the p110 PI3-kinases. Indeed, almost 20 years ago the PTEN-negative human glioblastoma cell line U251-MG was observed to decrease proliferation only when wild-type PTEN was expressed in the nucleus, while cytoplasmic PTEN had no effect on growth of these human cells (24). Further suggesting a novel mechanism operating in these cells, nuclear PTEN expression had no effect on AKT phosphorylation, suggesting the nuclear PTEN pathway is de-coupled from PI3-kinase signaling (24). Although not the only way PTEN operates in cancer, these data support the development of inhibitors against IPMK, which could be effective in halting growth of PTEN-negative tumors, particularly human glioblastomas.

Here, we report development of both 1^st^ and 2^nd^ generation IPMK inhibitors, characterizing their effects in U251-MG glioblastoma cells, and pharmacokinetic properties in mice. Compound 15 had the best pharmacokinetic properties, and treatment of U251-MG glioblastoma cells with this compound led to a novel metabolic profile which has not been previously observed in genetic loss-of-function studies of IPMK: compound **15** decreased only InsP_5_ levels, without effecting labeled InsP_6_ abundance in ^3^H-inositol metabolic labeling studies of human U251 glioblastoma cells. These data suggest chemical inhibitors of IPMK can reveal novel metabolic profiles in human glioblastoma cells not been previously observed using genetics. The compounds presented here will form the basis of future efforts to develop IPMK inhibitors with potential for clinical applications, particularly in human glioblastoma.

## RESULTS

### Compound 1 (UNC7437) regulates human glioblastoma cell line genes associated with cancer and viral infection/inflammation

In our efforts to develop IP6K1 inhibitors (25), we fortuitously discovered that compound **1** had off-target efficacy toward IPMK kinase activity. The specific inhibition of IPMK kinase activity was confirmed through extensive binding, activity and a co-crystal structure of human IPMK all reported elsewhere, which together suggested compound **1** selectively inhibits pure, recombinant human IPMK, when compared to the related human IP6K1 enzyme. Here, we report biological characterization compound **1** (**Fig 1A**), with an IC_50_ by Kinase-Glo assay of 26.2 ±1.2nM. Several previous studies have used genetic analyses to link the kinase activity of IPMK with gene expression and transcriptional regulation (6). We therefore asked if rapid treatment with compound **1** might regulate gene expression in a cell-based model of human cancer, the human glioblastoma cell line U251-MG (U251). The growth of U251 cells is halted by specific expression of nuclear PTEN, whereas cytoplasmic PTEN has no effect on growth of these cells. This observation is significant as IPMK can counteract the effects of PTEN, and expression of both PTEN and IPMK is often observed in the nucleus. Further, the growth effects in U251 cells were independent of any changes to AKT phosphorylation, and constitutively active AKT could not rescue the effects, suggesting PTEN acts through a PI3-kinase independent mechanism in U251 cells (24). Thus, we used U251 cells to test the efficacy of the first-generation compound **1**. Treatment with 400nM compound **1** (30X over the IC_50_) resulted in a significant decrease in cell number as determined by CyQuant DNA content assay (**Fig 1B**), suggesting a decrease in proliferation of U251 glioblastoma cells was induced by compound **1**. Identical treatment of these cells significantly decreased accumulation of InsP_4_, InsP_5_ and InsP_6_, as measured by ^3^H-inositol labeling followed by HPLC (**Fig 1C-D**). These data suggest compound 1 decreased cell growth and the expected inositol phosphates in human U251-MG cells.

**Figure 1:**
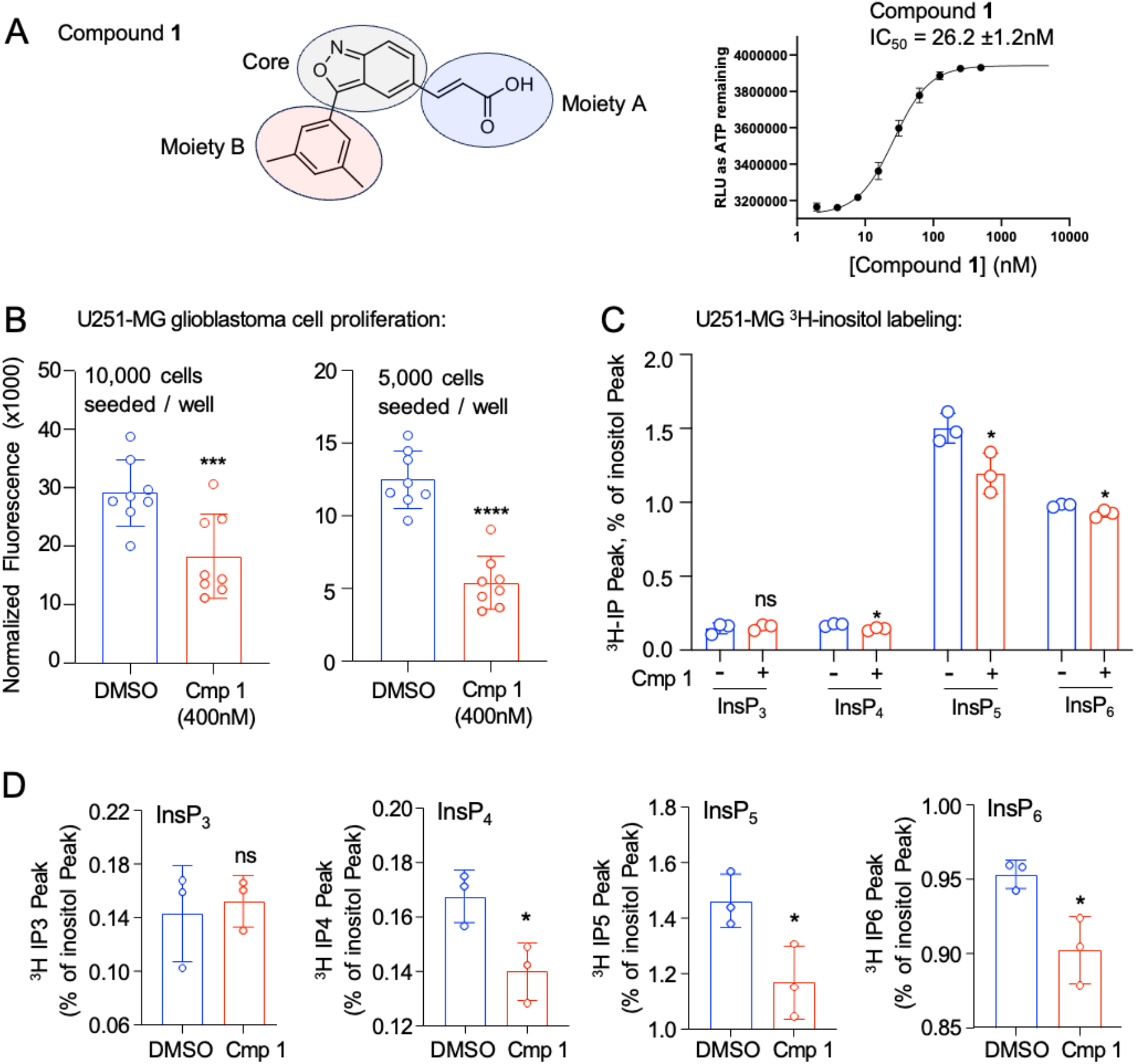
*Compound 1 inhibits human U251 glioblastoma cell growth and labeled inositol phosphate levels*. **A**. Chemical structure of compound 1, indicating moiety A (blue highlight), the core (gray highlight) and moiety B (red highlight), and the IC_50_ value as determined by kinase-GLO assay. **B**. CyQuant cell count assay of 10,000 human U251-MG glioblastoma cells seeded per well, treated 48 hours with DMSO vehicle or 400nM compound 1 (n=8, ***p=0.0046, unpaired t-test). **C**. 400nM compound 1 treatment for 48 hours of 3H-inositol metabolically labeled U251-MG cells, analyzed by HPLC, expressed as the percentage of the inositol peak from the HPLC for lnsP3, lnsP4, lnsP5 and lnsP6, as indicated, *p<0.05 by unpaired t-test Bonferroni corrected for multiple comparisons. **D**. Same data as in C re-plotted with a non-zero y-axis to facilitate visualizing differences between DMSO vs. compound treatment for each indicated inositol phosphate species, *p<0.05 by unpaired t-test Bonferroni corrected for multiple comparisons. *These data suggest compound **1** decreases U251 cell growth and inositol phosphate levels, consistent with inhibiting IPMK kinase activity in vivo*.

### Compound 1 regulates genes important for cell growth and inflammatory responses

IPMK and inositol phosphates have been linked to transcriptional regulation (21, 26–30), so we next performed RNA-seq transcriptome analysis of these cells after 400nM compound **1** *vs*. DMSO control for 48 hours (**Fig 2A**). These transcriptome analyses identified 993 transcripts differentially expressed (*p*_*adj*_<0.05) that also had a log2 fold change (log_2_FC) > ±1. Among these genes, 661 transcripts were downregulated while 332 were upregulated. We subjected the differentially expressed transcripts to gene set enrichment analysis (GSEA) using the Hallmark Pathways collection from the molecular signatures database (**Fig 2B**), showing several pathways associated with cancer or tumor metastasis were significantly associated with the gene sets regulated by compound **1** treatment (apical junction, mitotic spindle, epithelial-mesenchymal transition (EMT), MYC targets, KRAS signaling). A similar GSEA analysis using all gene sets in molecular signatures identified enrichment of genes belonging to two categories: 1) cancer gene sets and 2) viral / inflammation gene sets (**Supplemental Fig S1**), although other gene sets were also significantly enriched as well, see supplemental spreadsheet for the full list of all enriched gene sets. Compound **1**-regulated transcripts were enriched in the Verhaak-Glioblastoma Classical gene set (**Fig 2C**) and in NFKBIA target genes (**Supplemental Fig S1**), while compound **1**-regulated transcripts were significantly de-enriched in several virus, vaccine and apoptosis gene sets (**Fig 2D, Supplemental Figure S1**). Together, these data suggest compound **1** decreases proliferation of these human U251 glioblastoma cancer cells and alters expression of genes enriched in cancer and inflammatory pathways.

**Figure 2.**
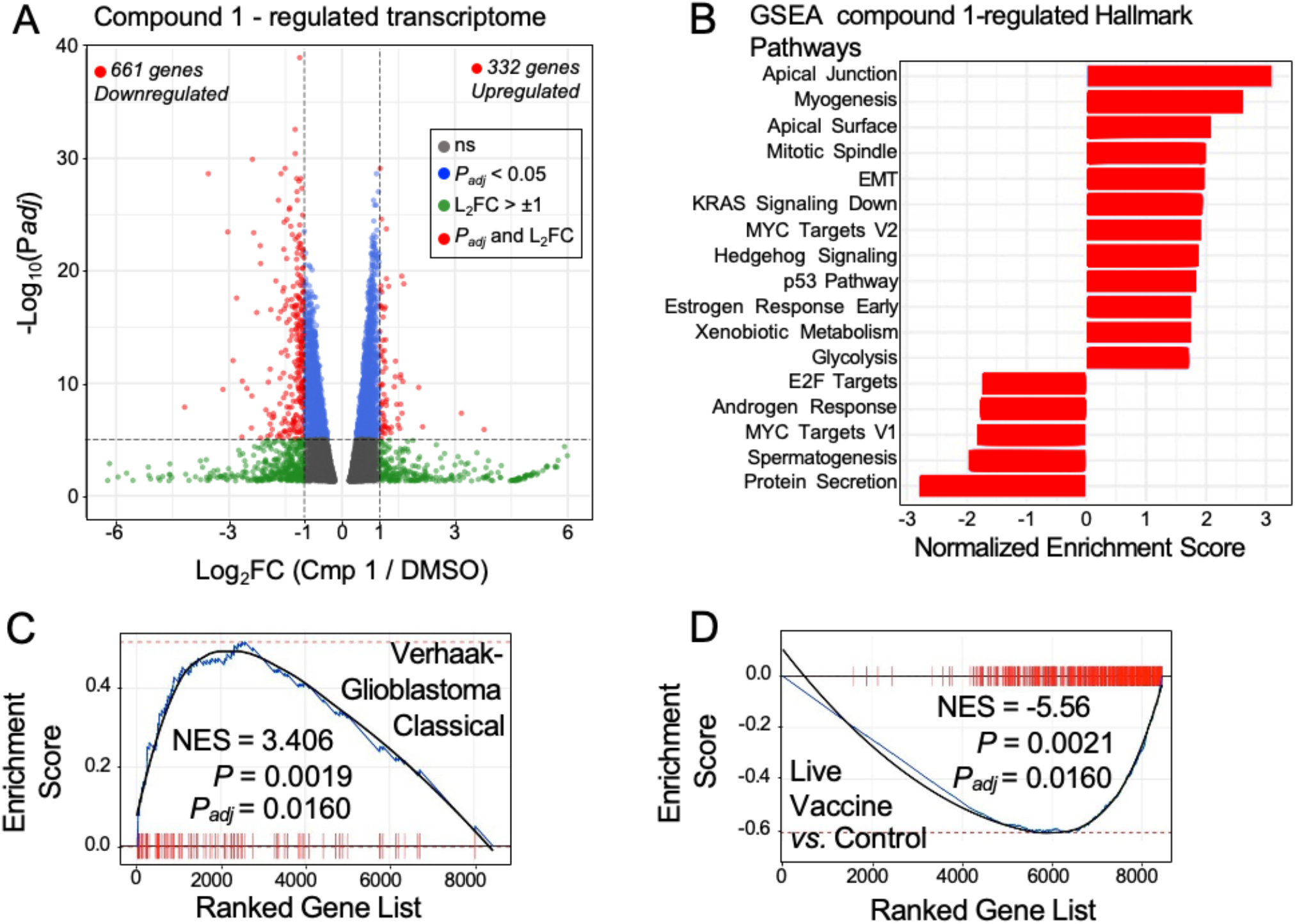
*Compound 1 inhibits cell growth and inositol phosphate abundance, while inducing a in human U251-MG glioblastoma cells*. **A**. Volcano plot from biological triplicate RNA-seq of human U251-MG glioblastoma cells, showing differentially expressed transcripts upon treatment with 400nM compound 1 for 48 hours, compared to DMSO vehicle control. **B**. Gene set enrichment analysis (GSEA) of the compound 1-differentially expressed gene set showing enrichment in several hallmark cancer pathways. **C**. Bar code GSEA plots from representative enriched and **D**. de-enriched gene sets using the Molecular Signatures database. *These data suggest compound 1 treatment of human glioblastoma cells regulates gene sets associated with cancer and viral infection/inflammation, consistent with the known functions of IPMK and inositol phosphates*.

### Modification of Moiety A to improve potency and drug-like properties

We synthesized carboxylic acid derivatives **2** and **3** (**Fig 3A-B**), compound **4** was the precursor for compound **3**. A Corey-Chaykovsky cyclopropanation of **4** with sulfonium ylide formed *in situ* from trimethylsulfoxonium iodide followed by an ester hydrolysis reaction under the basic conditions yielded the desired analog **2**. Synthesis of **4** is straightforward since both starting materials 2-phenylacetonitrile and 5-(4-nitrophenyl)-1*H*-tetrazole are commercially available. A ring formation reaction between 2-phenylacetonitrile and 5-(4-nitrophenyl)-1*H*-tetrazole under the basic condition yielded the desired compound **4** in 26% yield. This cyclization reaction is also the key step for other tetrazole analog synthesis and can be performed in a multigram scale.

**Figure 3.**
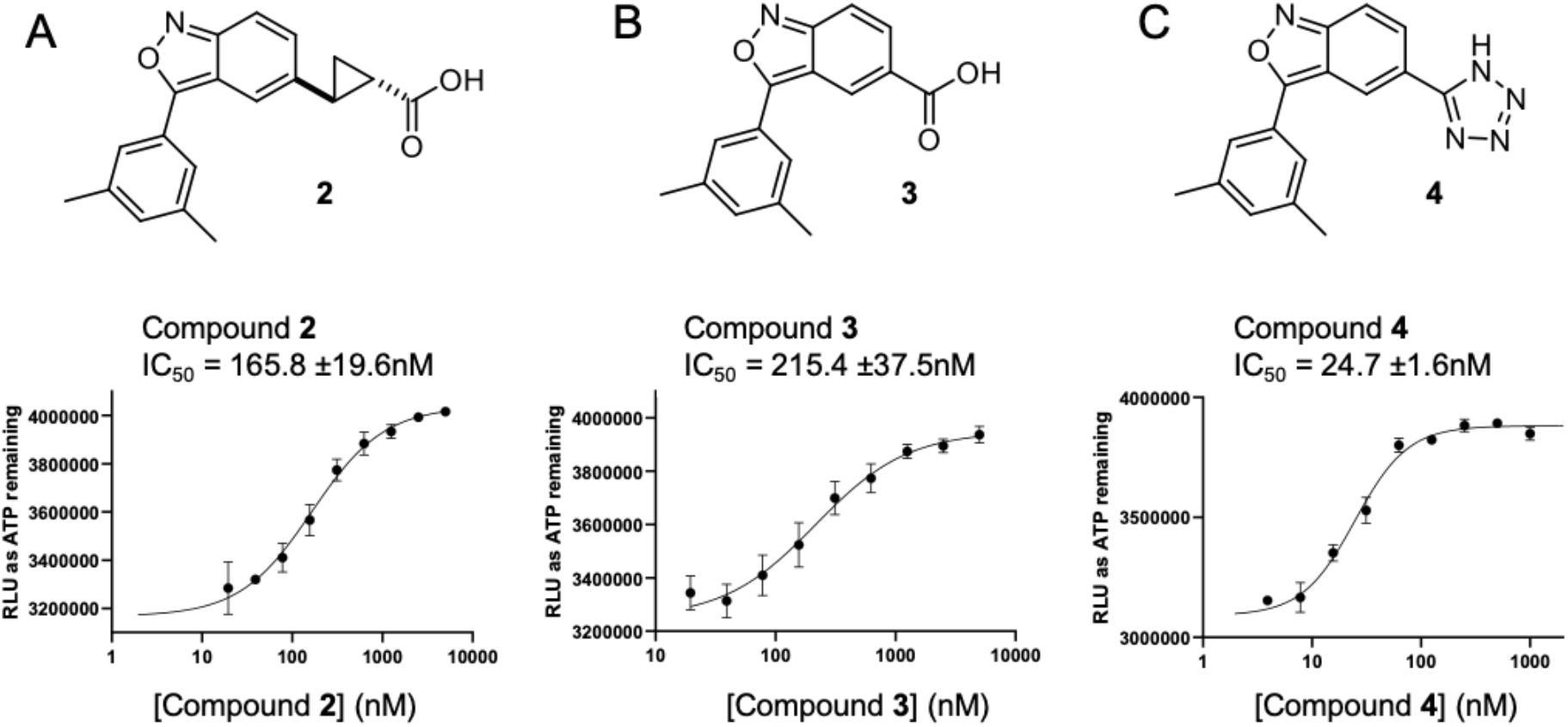
Chemical structures, IC_50_ values and curve fits for **A**. compound **2**, **B**. compound **3** and **C**. compound **4**.

### Modification of Moiety A to improve potency and drug-like properties

The double bond on the acrylic acid side chain in 1 may bring liability to be a potential Michael acceptor.

We synthesized Compound **2** via the route shown in **Scheme 1**. We Therefore, we replaced the double bond with a cyclopropyl group to yield compound **2** (**Fig 3A-B**) which contains two stereocenters that would likely necessitate separation into pure enantiomers (**Fig 3A**). On the other hand, we truncated the acrylic acid side chain to an acetyl group to produce analog **3** (**Fig 3B**), which is expected to have decreased potency for IPMK as previous studies have shown that related inhibitors with carboxylic acid modifications at this position show decreased selectivity towards IPMK(25). The carboxylic acid functional group in Compound **3** often leads to lower permeability to biological membranes, thus is frequently replaced by an acid bioisostere.^2-3^ In our effort to identify a suitable acid bioisostere, we found compound **4**, containing a tetrazole, displayed improved solubility (**Fig 3C**). The syntheses of compounds **2 & 4** are presented in Scheme 1. Compound **5** was the precursor for compound **2** (**Scheme 1A**). A Corey-Chaykovsky cyclopropanation of **5** with sulfonium ylide formed *in situ* from trimethylsulfoxonium iodide followed by an ester hydrolysis reaction under the basic conditions yielded the desired analog **2**. Synthesis of **4** is straightforward since both starting materials, 2-phenylacetonitrile and 5-(4-nitrophenyl)-1*H*-tetrazole, are commercially available (**Scheme 1B**). A ring formation reaction between 2-phenylacetonitrile and 5-(4-nitrophenyl)-1*H*-tetrazole under the basic condition yielded the desired compound **4** with a 26% yield. This cyclization reaction is also the key step for other tetrazole analog synthesis and can be performed on a multigram scale.

**Scheme 1.**
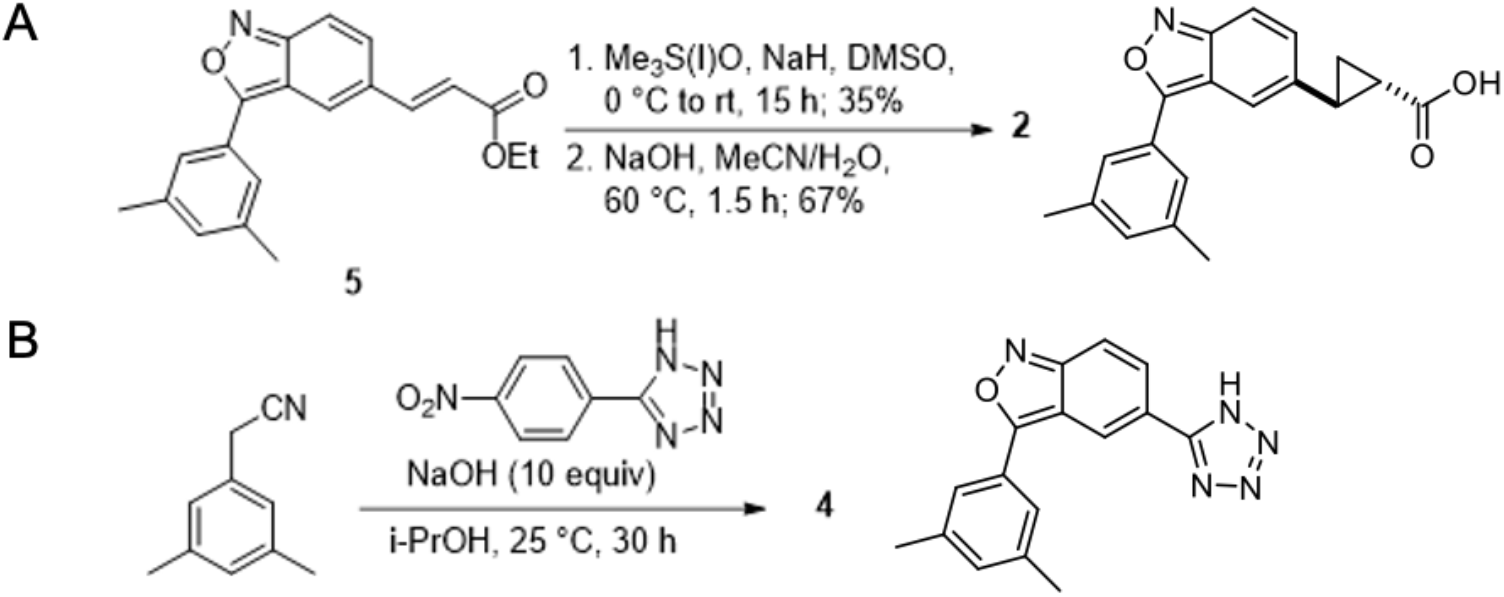
Analog Synthesis of **A**. compound **2**. and **B**. compound **4**.

Our next focus was to change moiety B to improve the pharmacokinetic profile of these compounds (**Fig 4A**). Compound **6** (**Fig 4B**) was produced by removal of both methyl groups on the phenyl ring from compound **4**. Compounds **7** (**Fig 4C**) and **8** (**Fig 4D**) were synthesized with smaller trifluoromethyl and trifluoromethoxy groups at the *para*-position of the phenyl. Compound **9** (**Fig 4E**) had a large phenoxy group at this position. We then synthesized compound **10** (**Fig 5B**), which bears a cyclopropanecarboxamide group. Compound **11** (**Fig 5C**) moved the cyclopropanecarboxamide group to the *meta*-position of the phenyl ring.

**Figure 4.**
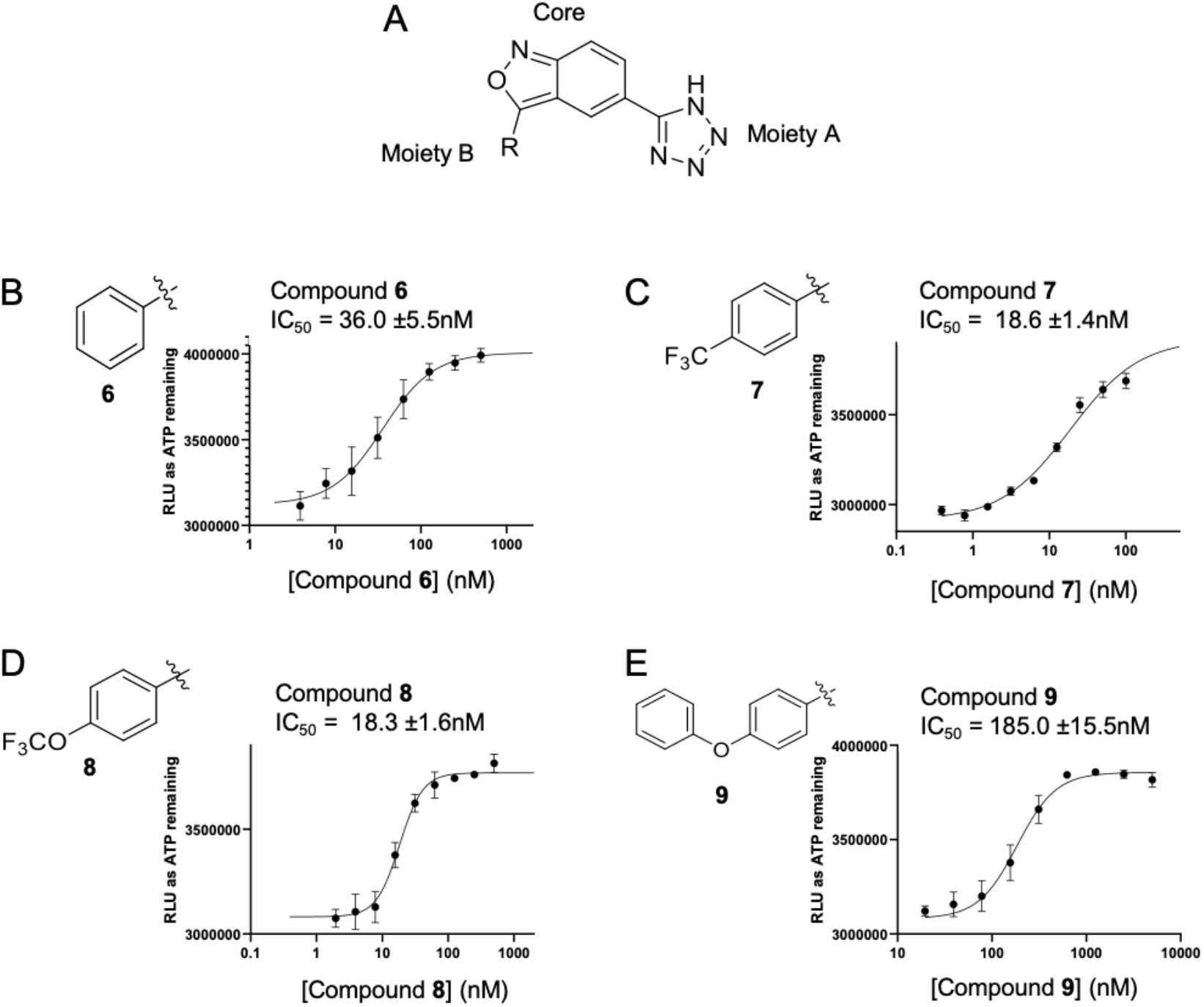
Chemical structure of the **A**. backbone scaffold of compound **4** with moiety **B** as the R group and moiety A and the core indicated. Chemical structures of the R groups (moiety B) for **B**. compound **6, C**. compound **7, D**. compound **8** and **E**. compound **9. F**.

**Figure 5.**
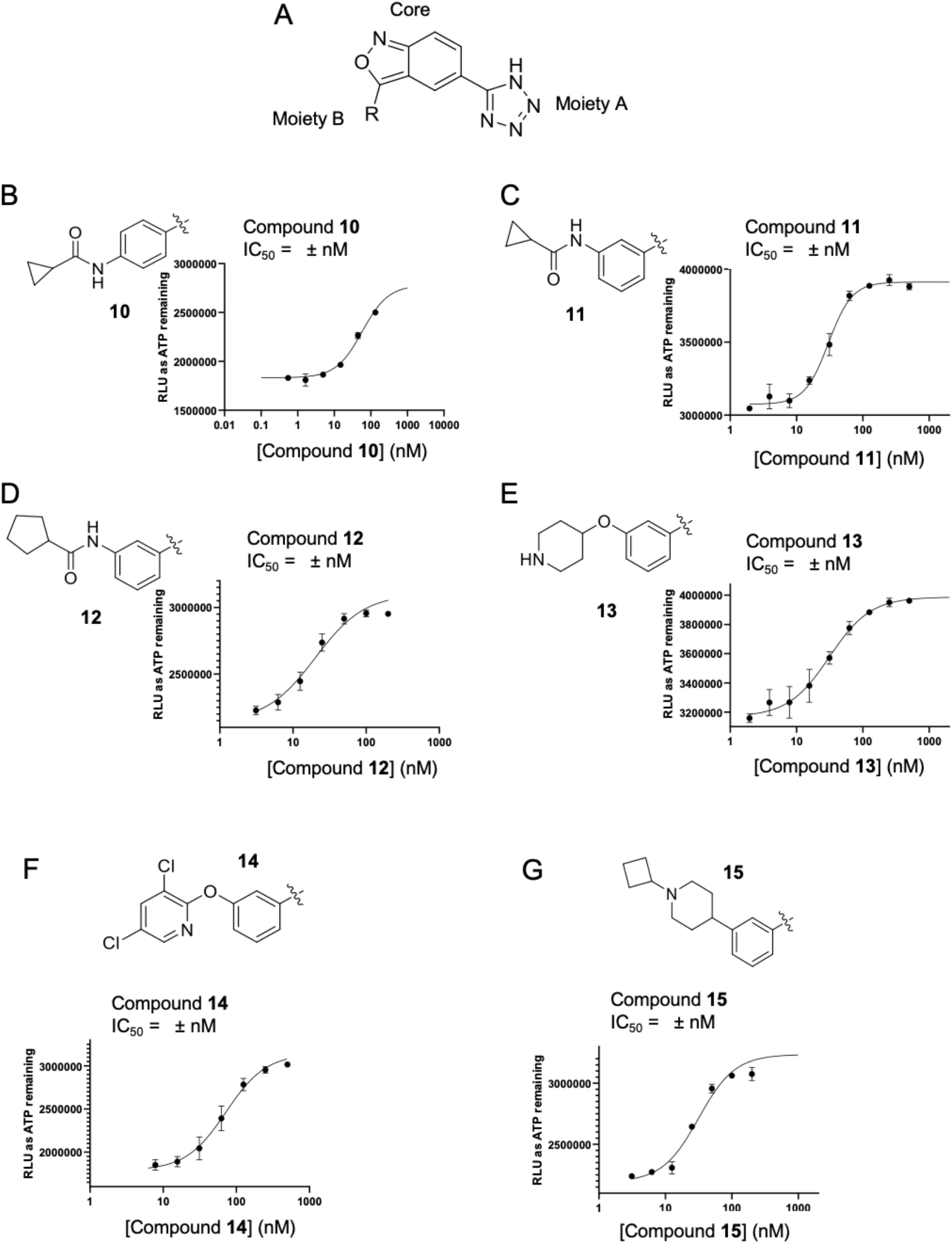
Chemical structure of the **A**. backbone scaffold of compound **4** with moiety B as the R group and moiety A and the core indicated. Chemical structures of the R groups (moiety B) for **B**. compound **10, C**. compound **11, D**. compound **12, E**. compound **13, F**. compound **14** and **G**. compound **15**.

Accordingly, we then examined the pharmacokinetic properties (PK) of compound **11**, in mice via intravenous (iv) administration with 3.0 mg/kg dose (**Table 1**). Compound **11** had a clearance (CL) of 7.5 mL/min/kg, but a short half-life of just over 1 hour (T_1/2_ 1.09 h) due to low volume of distribution (Vss, 0.21L/kg, less than mouse blood volume of 0.70 L/kg)^4^. To increase the volume of distribution, we next attempted to introduce basic groups into these inhibitors.

**Table 1.** In Vivo Pharmacokinetic Parameters of selected analogues (3 mg/kg, n = 2 mice per time point)

| Compound<sup>a</sup> | $T_{1/2}$<br>(h) | Cmax<br>(μM) | AUClast<br>(h*μM) | Vss<br>(L/kg) | CL<br>(mL/min/kg) |
| --- | --- | --- | --- | --- | --- |
| **11** | 1.09 | 66.6 | 19.4 | 0.21 | 7.52 |
| **13** | 1.29 | 44.0 | 10.8 | 0.23 | 11.8 |
| **14** | 2.01 | 50.5 | 7.7 | 0.27 | 15.5 |
| **15<sup>b</sup>** | 1.95 | 28.3 | 4.6 | 0.39 | 22.8 |
| **15<sup>c</sup>** | 1.36 | 52.3 | 10.1 | 0.71 | 37.9 |

<sup>a</sup>Intravenous formulation: 10% NMP, 5% solution in normal saline (0.9% NaCl). <sup>b</sup>Dose: 2.75mg/kg, <sup>c</sup>dose: 10mg/kg, n=3.

Compounds **13-15** (**Fig 5E-G**) incorporated basic groups such as 4-piperidine and 3,5-dichloro-pyridine at the meta-position of the phenyl ring. Compound **13** (**Fig 5E**) had a 4-piperidinyloxy group at the phenyl ring while compound **14** (**Fig 5F**) contains a meta-3,5-group, and the volume of distribution was increased for all three compounds **13-15** (**Table 1**). Compound **15** had a larger volume of distribution (0.39L/kg) and longer half-life (1.95h) while retaining low clearance (22.8mL/min/kg). This was further evaluated in mice with a 10 mg/kg dose via intravenous and intraperitoneal administration (n=3). Compound **15** had large AUClast (8.26h*µM) and 82% bioavailability. Thus, compound **15** displays excellent *in vivo* pharmacokinetics, suggesting compound **15** has the potential for *in vivo* utility.

We then determined the IC_50_ values for select compounds against IPMK to determine if these modifications improved potency (**Table 2**). The most highly potent molecule was compound **12** (20.1 ±2.0nM.). However, compound **15** had comparable potency (31.6 ±2.4nM) and excellent *in vivo* pharmacokinetics, suggesting it as the best candidate to pursue using additional cell-based assays.

**Table 2.**
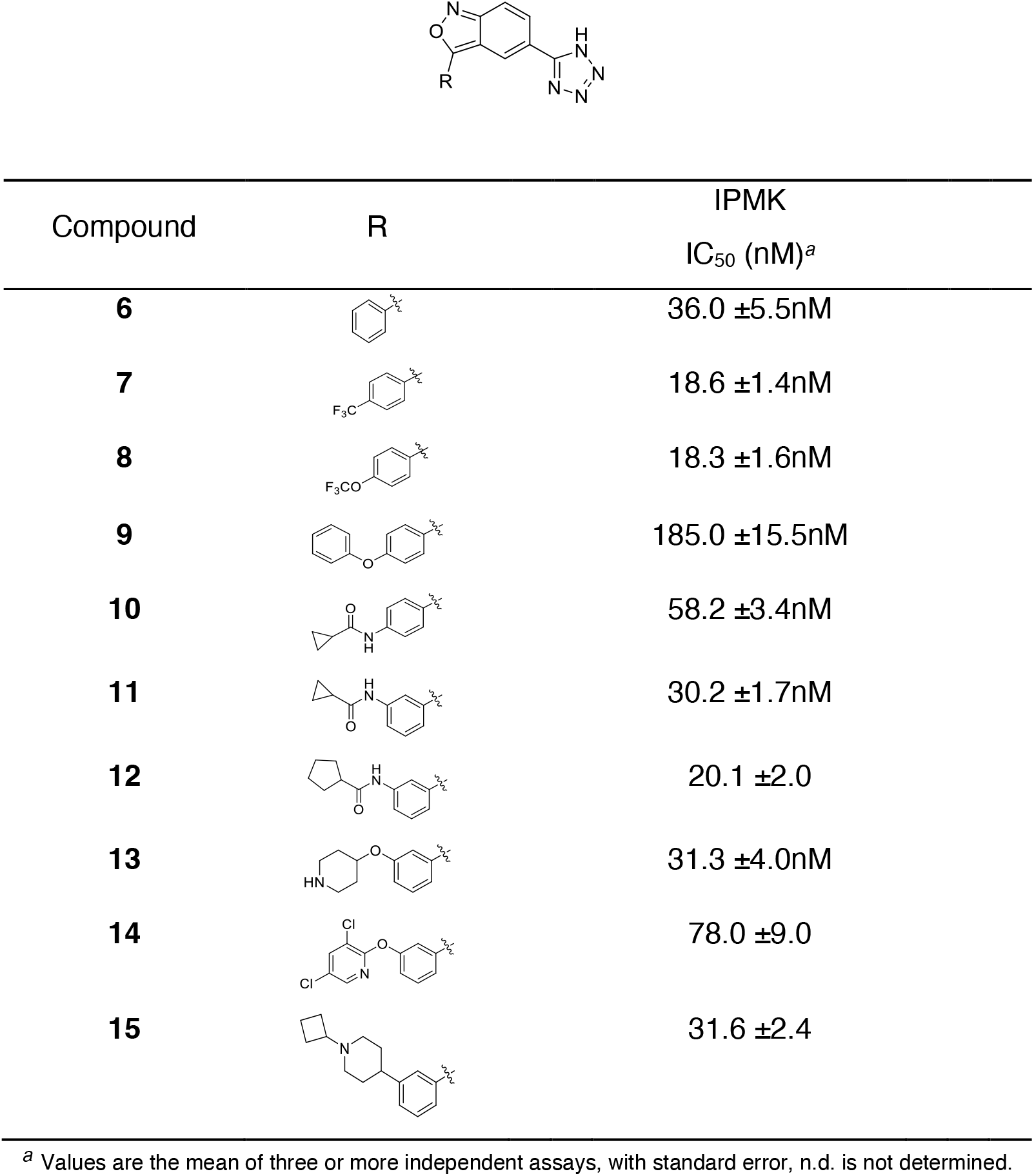
Modification of Moiety B to increase selectivity.

**Table 3:** Compound Numbering Chart.

| Compound | UNC numbering | IPMK<br>IC<sub>50</sub> (nM)<sup>a</sup> |
| --- | --- | --- |
| **1** | UNC7437A | 26.2 ±1.2nM |
| **2** | UNC7789A | 165.8 ±19.6nM |
| **3** | UNC7438A | 215.4 ±37.5nM |
| **4** | UNC7854A | 24.7 ±1.6nM |
| **6** | UNC7782A | 36.0 ±5.5nM |
| **7** | UNC7855A | 18.6 ±1.4nM |
| **8** | UNC7891A | 18.3 ±1.6nM |
| **9** | UNC7890A | 185.0 ±15.5nM |
| **10** | UNC8787A | 58.2 ±3.4nM |
| **11** | UNC9021A | 30.2 ±1.7nM |
| **12** | UNC9022A | 20.1 ±2.0 |
| **13** | UNC9120A | 31.3 ±4.0nM |
| **14** | UNC9107A | 78.0 ±9.0 |
| **15** | UNC9750A | 31.6 ±2.4 |

<sup>a</sup> Values are the mean of three or more independent assays, with standard error

### Compound 15 decreases human glioblastoma cel growth and selectively decreased cellular InsP_5_ abundance

We next tested the effect of compound **15** on various functional aspects of the human glioblastoma cancer cell line U251-MG. Compound **15** significantly decreased cellular growth (1.0μM, 48 hours), with a larger effect size occurring at a higher dose of Compound 15 (10μM, 48 hours, **Fig 6A**). Using tritiated, ^3^H-inositol metabolically labeled cells treated with 10μM compound **15** for 48 hours, InsP_4_ levels increased significantly, while InsP_5_ levels decreased significantly. However, no change was detected in the level of InsP_6_ (**Fig 6B-D**). This unexpected result contrasts with results obtained from gene deletion studies involving the loss of the *IPMK* gene, where both InsP_5_ and InsP_6_ decreased to undetectable levels (20). Complete IPMK gene loss requires several weeks of cellular selection to obtain the knockout, whereas chemical inhibition of the IPMK happens within minutes to hours, highlighting the differences between chemical inhibition vs. complete genetic loss, which provides a potential explanation for these differences. We note that while InsP_4_ can be generated by other enzymes in mammalian cells, the only known source of InsP_5_ in mammalian cells is IPMK.

**Figure 6.**
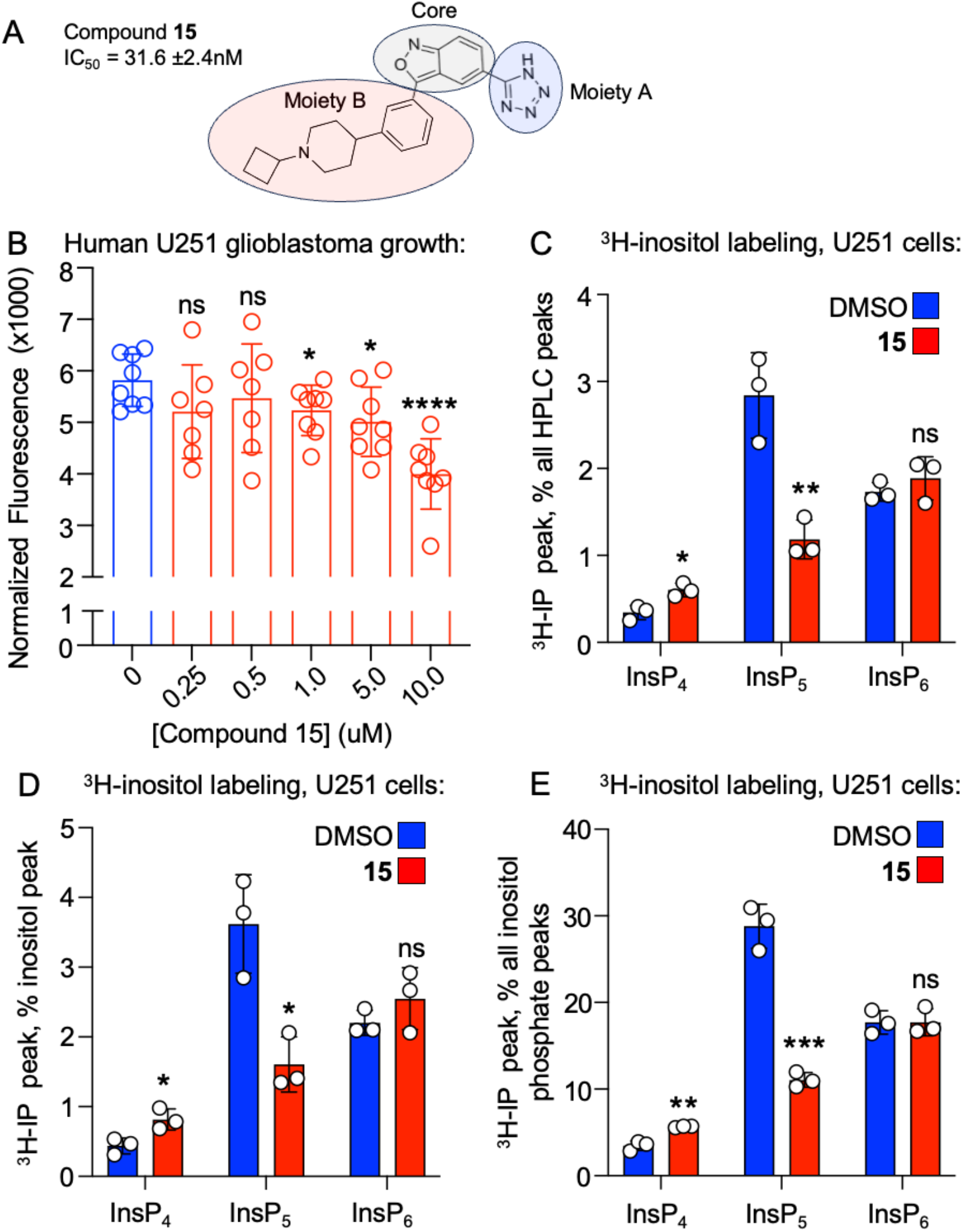
*Compound 15 inhibits cell growth and lnsP*_*5*_ *abundance in human U251-MG glioblastoma cells*. **A**. Chemical structure of **15**, error for IC_50_ for IPMK represents standard error from 3 independent assays. **B**. CyQuant cellular proliferation assay of 10,000 human U251-MG glioblastoma cells seeded per well, treated 48 hours with DMSO vehicle or 10uM compound 15 (n=8, *\*\*\*Padf0*.*0001* by one way ANOVA Dunnett’s corrected, *\*p<O*.*05* by unpaired t-test). **B**. 48 hours of 10uM Compound 15 treatment was used to treat ^3^H-inositol metabolically labeled U251-MG cells, analyzed by HPLC, expressed as a percentage of all the peaks detectable from the HPLC, *\*p<0*.*05, **p<0*.*01, ***p<0*.*001* by unpaired t-test compared to DMSO treated control for all remaining panels, n=3. **C**. Same as B but expressed as percentage of the inositol HPLC peak, n=3. **D**. Same as C but expressed as a percentage of all the inositol phosphate peaks combined from the HPLC, n=3. *These data suggest 48 hours of compound 15 treatment decreases labeled lnsP*_*5*_, *but does not change detectable labeled lnsP*_*6*_ *levels, in U251-MG human glioblastoma cells*.

Together, these data suggest that 48 hours treatment with compound **15** not only inhibited human U251-MG glioblastoma cell growth, but also induced a novel, specific decrease in cellular InsP_5_, a direct product of IPMK kinase activity, while having no detectable effect on InsP_6_. Together, the data suggest that rapid (48hr) treatment with compound **15**, selectively and effectively inhibits InsP_5_ levels, without affecting InsP_6_. To our knowledge, the observed decrease in InsP_5_ with no change to InsP_6_ has not been previously observed in any genetic knockout studies of IPMK in human cells. Together, the data suggest that these new compounds have potential clinical utility for testing as a treatment in glioblastoma, and as a tool to further basic science research investigating inositol phosphate and IPMK signaling, for the first time independently of the side effects inherent to long-term genetic knockout studies.

## DISCUSSION

This study suggests that small molecule IPMK-selective inhibitors may have therapeutic value in slowing cancer cell growth, particularly in glioblastoma. The U251-MG cells used for cellular characterization of the compounds developed in this study were selected because they are PTEN-negative, and a previous study from Alfred Yung’s lab had shown that proliferation of these cells can be only be reduced by the expression of wild-type PTEN and not a Cytoplasmic expression of PTEN had no effect on growth of these U251-MG cells (24). Attenuation of proliferation by nuclear, wild-type PTEN was independent of any effects on AKT-phosphorylation, suggesting AKT activation by classic PI3-kinase signaling may play a less important role in human glioblastoma U251 cell growth. Our data showing that several IPMK kinase inhibitors reduced U251-MG cell growth represents a pharmacological recapitulation of the slow-growth phenotype induced by overexpression of nuclear PTEN in these same human U251-MG glioblastoma cells. It is therefore tempting to speculate that chemical inhibitors of IPMK may be capable of uniquely targeting PTEN-negative glioblastoma, particularly in patients that have not responded well to therapies that decrease AKT phosphorylation. Indeed, further improvements to compound **15** based on structure-activity relationship studies, deserves thorough consideration as a potential future therapy in PTEN-negative glioblastoma or other brain cancers.

The purified human IPMK enzyme can generate both InsP_4_ and InsP_5_. In cells, InsP_4_ can also be generated by IP3K, but InsP_5_ production is only known to occur through IPMK in human cells. IPMK-generated InsP_5_ is the required precursor for generation of InsP_6_ by IPK1. For this reason, when the IPMK gene has been genetically removed from cells, those cells have decreased labeled InsP_5_ and concomitantly decreased labeled InsP_6_. Compound **15** decreased levels of InsP_5_ but had no effect on InsP_6_ after 48 hours of treatment, a novel observation that highlights the importance of orthogonal chemical validation of genetic results. These data are consistent with compound **15** rapidly attenuating IPMK activity in the U251 cells with high specificity, which prevents label incorporation into the most direct product of IPMK kinase activity (InsP_5_).

It is also unclear why InsP_4_ levels in glioblastoma cells increased with compound **15** treatment yet decreased with compound **1** treatment. Here we note that InsP_4_ levels also increase upon genetic deletion of IPMK in mammalian HT-29 cells (20). A reasonable hypothesis explaining this biological phenomenon is that compound **1** may decrease InsP_4_ due to off-target effects on IP3K and/or ITPK1 enzymes, which both generate InsP_4_. Furthermore, upregulation of InsP_4_ by compound **15** could be due to compensatory biological upregulation of IP3K and/or ITPK1 and consequent increased production of Ins(1,3,4,5)P_4_ and Ins(1,3,4,6)P_4_, respectively, preventing InsP_4_ decreases due to IPMK inhibition. Disentangling the complex network of inositol phosphate biological regulation will be important in future studies, expedited by the new series of compounds reported here. Clinically, these studies also put forth the possibility of inhibiting IPMK as a therapy in PTEN-negative glioblastoma multiforme, given the favorable pharmacokinetic profiles of this series of kinase inhibitors. Together, these studies introduce the complete synthesis and initial biological characterization of a series of 1^st^ and 2^nd^ generation IPMK-directed inhibitors, which now form the basis for further medicinal chemistry optimization for eventual movement into the clinic.

## ACKNOWLEDGMENTS

The authors thank Dr. Stephen Shears and his team at NIEHS for helpful discussions. This work was supported by the University of North Carolina Cancer Research Fund to X.W., National Institutes of General Medical Sciences R01 GM132592 to R.D.B, and NIA AG071975 to L.E.R. We thank Dr. Zibo Li and Dr. Xuedan Wu for their help in the HPLC data collection for purity determination of key compounds.

## ABBREVIATIONS USED

InsP_3_,: inositol 1,4,5-trisphosphate
InsP_4_,: inositol tetraphosphate
InsP_5_,: inositol pentakisphosphate
InsP_6_,: inositol hexakisphosphate
ip,: intraperitoneal
IP6K,: inositol hexakisphosphate kinase
iv,: intravenous

## MATERIALS & METHODS

### ^3^H-inositol metabolic labeling and HPLC

Human U251-MG glioblastoma cells (ECACC 09063001) were grown in DMEM supplemented with 10% FBS. The medium was changed to inositol-free DMEM and cells were inositol-starved for 1-3 days. Cells were maintained in inositol-free DMEM medium and supplemented with 100µCi ^3^H-inositol (Perkin Elmer) for 2 days. Cells were then washed in PBS and lysed in 200µL HCl (0.5M), 250µL(31) of MeOH to one well of a 6-well plate (approximately 10^6^ cells), followed by 125µL 2M KCl. Lysates were scrapped and sonicated in ice for 3 one-minute cycles. Inositol phosphates (IPs) extracts were cleaned of lipids by adding 250μl of CHCL_3_, vortexing and centrifuging samples for 5 minutes at 14,000 rpm to separate upper aqueous and lower organic phases, upper phase contains soluble IPs. This extraction was repeated once. Samples were dried in a speedvac, resuspended in 10 mM ammonium phosphate (pH= 3.5) and filtered. HPLC separation and detection of the inositol species on the upper phase used a Partisphere SAX 4.6 x 125 mm column, eluted with 10mM (Buffer A) to 1.7M (buffer B) ammonium phosphate (pH= 3.5) and on-line flow scintillation analyzer (FSA, Perkin-Elmer). Elution of [^32^P]-labeled IP4, IP5 and IP6 standards confirmed the identity of each peak off the HPLC. Quantification represents three biologically independent metabolic labeling samples with error representing standard deviation of the mean.

### RNA-seq and gene set enrichment analysis

DNase-treated RNA prepped from one well of a six well plate (approximately 10^6^ cells) was extracted using Quick-RNA MiniPrep (Zymo Research) from human U251-MG glioblastoma cells (ECACC 09063001) after 48 hours treatment with Compound 1. PolyA selected RNA-seq libraries were prepared and sequenced at the Vanderbilt Technologies for Advanced Genomics (VANTAGE) core on an Illumina NextSeq500, bases and reads with low quality scores or reads, respectively, were removed and Illumina library adapter sequences were trimmed using bbDuk (https://jgi.doe.gov/data-and-tools/software-tools/bbtools/bb-tools-user-guide/bbduk-guide/). The resulting files were mapped to hg38 using HiSat2(32), poorly mapped reads were removed with SamTools(33). Reads mapping to RefSeq genes were counted using HTSeq-count(34) and differential expression assessed using DESeq2(35) with Apeglm LFCShrinkage(36). Gene sets from the molecular signatures database (version 6.1) were used for Gene Seta Enrichment Analysis (GSEA), those data provided as supplemental spreadsheets, all original FASTQ files and underlying count data are available upon request.

### IC50 values determinations

IC50 values were determined using the purified recombinant human IPMK kinase domain expressed in *E*.*coli* and Kinasethe Kinase-Glo assay, as previously described(2, 37, 38). This human IPMK kinase domain (130 nM) was used in reaction buffer (20 mM HEPES pH 6.8, 100 mM NaCl, 6 mM MgCl_2_, 20 µg/mL BSA, 1mM DTT)containing 12.5µM Ins(1,4,5)P3 kinase substrate at room temperature, and the reactions were analyzed after 12 minutes. These assay conditions were optimized with respect to reaction time, IPMK enzyme concentration and IP_3_ concentration determine the IC50 for these ATP-competitive compounds.

### Pharmacokinetics

All the procedures related to animal handling, care, and the treatment in this study were performed according to guidelines approved by the Institutional Animal Care and Use Committee (IACUC) of Pharmaron (PK-R-06012023 and PK-M-07182023) following the guidance of the Association for Assessment and Accreditation of Laboratory Animal Care (AAALAC). A group of 3 male CD1 mice (2 for exploratory PK) were dosed intravenously (iv) or intraperitoneal (ip) with solution formulation of tested compounds in 10% NMP, 5% Solutol in normal saline (v/v/v, 10:5:85). From each mouse, blood samples (30 mL/sample) were collected from orbit vein such that samples were obtained at 0.083, 0.25, 0.5, 1, 2, 4, 8 & 24 post dose. At each time point blood samples were collected from three mice in labeled micro centrifuge tube containing EDTA-K2 as anticoagulant. Plasma samples were separated by centrifugation of whole blood and stored at −75±15ºC until bioanalysis. All samples were processed for analysis by precipitation using acetonitrile (ACN) and analyzed with fit for purpose LC/MS/MS method (Below quantifiable limit (BLOQ) was 5.0 ng/mL). Pharmacokinetic parameters were calculated by non-compartmental analysis model using WinNonlin 8.3.

### Graphical software

Graphs have been prepared using SigmaPlot v14 and GraphPad Prism 9; IC_50_ data were calculated using GraphPad Prism 9.

## Supplementary Information

## ABBREVIATIONS USED

[Bmim]BF_4_: 1-butyl-3-methylimidazolium tetrafluoroborate
*t*-BuOH: *tert*-butanol
DIO: diet-induced obesity
DMEM: Dulbecco’s modified eagle medium
dpm: disintegrations per minute
EWAT: epididymal-adipose tissue
FBS: fetal bovine serum
GTT: glucose tolerance test
HATU: 1-[bis(dimethylamino)methylene]-1*H*-1,2,3-triazolo[4,5-*b*]pyridinium 3-oxid hexafluorophosphate
HFD: high fat diet
HPLC: high-performance liquid chromatography
Ins(1,4,5)P_3_: inositol 1,4,5-trisphosphate
InsP_5_: inositol pentakisphosphate
InsP_6_: inositol hexakisphosphate
5-InsP_7_: 5-diphosphoinositol pentakisphosphate
ip: intraperitoneal
IP6K: inositol hexakisphosphate kinase
ITC: isothermal titration calorimetry
ITT: insulin tolerance test
iv: intravenous
IWAT: inguinal-adipose tissue
Me_4_tBuXPhos: 2-di-*tert*-butylphosphino-3,4,5,6-tetramethyl-2′,4′,6′-triisopropyl-1,1′-biphenyl
MeCN: acetonitrile
NAFLD: nonalcoholic fatty liver disease
NASH: nonalcoholic steatohepatitis
PP-InsP: inositol pyrophosphate
*i*-PrOH: isopropyl alcohol
PTT: pyruvate tolerance test
RuPhos: 2-dicyclohexylphosphino-2′,6′-diisopropoxybiphenyl
RuPhos-Pd-G2: chloro(2-dicyclohexylphosphino-2′,6′-diisopropoxy-1,1′-biphenyl)[2-(2′-amino-1,1′-biphenyl)]palladium(II)
SPhos: 2-dicyclohexylphosphino-2′,6′-dimethoxybiphenyl
T2DM: type 2 diabetes mellitus
TNP: *N*2-(*m*-Trifluorobenzyl)-*N*6-(*p*-nitrobenzyl)purine.

---

**Supplemental Figure S1.**
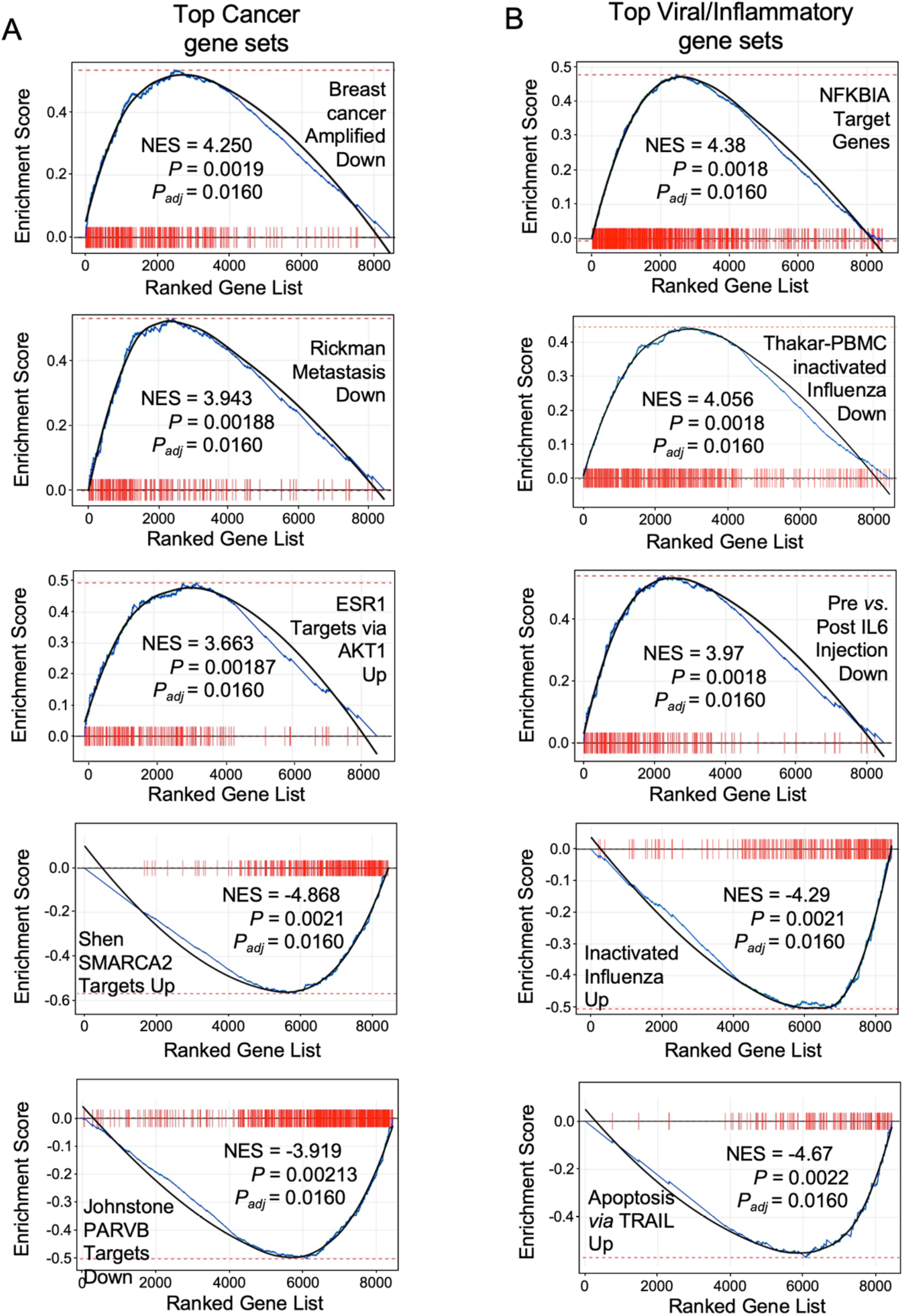
*Gene sets enriched in Compound 1-regulated transcripts*. Bar code GSEA plots from **A**. the top cancer-related gene sets and **B**. the top viral/inflammatory gene sets from the Molecular Signatures database. Normalized enrichment scores (NES) and *p*-values (both unadjusted and adjusted) are indicated in each enrichment plot.

## Synthesis of Compounds

Microwave reactions were carried out using a CEM Discover-S reactor with a vertically focused IR external temperature sensor and an Explorer 72 autosampler. The dynamic mode was used to set up the desired temperature and hold time with the following fixed parameters: PreStirring, 1 min; Pressure, 200 psi; Power, 200 W; PowerMax, off; Stirring, high. Sonication was carried out on Branson 3510 Ultrasonic Cell. Centrifugation was carried out on Eppendorf Centrifuge 5418. Flash chromatography was carried out on Teledyne ISCO Combi Flash^®^ R_f_ 200 with pre-packed silica gel disposable columns or pre-packed reverse phase C18 columns. Analytical thin-layer chromatography (TLC) was performed with silica gel 60 F_254_, 0.25 mm pre-coated TLC plates. TLC plates were visualized using UV_254_ and phosphomolybdic acid with charring. All ^1^H NMR spectra were obtained with a 400 or 500 MHz spectrometer using CDCl_3_ (7.26 ppm), DMSO-*d*^*6*^ (2.50 ppm, quintet) or CD_3_OD (3.31 ppm, quintet) as an internal reference. Signals are reported as m (multiplet), s (singlet), d (doublet), t (triplet), q (quartet), and bs (broad singlet); and coupling constants are reported in hertz (Hz). ^13^C NMR spectra were obtained with a 100 or 125 MHz spectrometer using CDCl_3_ (77.2 ppm, triplet), DMSO-*d*^*6*^ (39.5 ppm, septet), or CD_3_OD (49.3 ppm, septet) as the internal standard. LC/MS was performed using an analytical instrument with the UV detector set to 220 nm, 254 nm, and 280 nm, and a single quadrupole mass spectrometer using electrospray ionization (ESI) source.

Samples were injected (2 μL) onto a 4.6 x 50 mm, 1.8μM, C18 column at room temperature. A linear gradient from 10% to 100% B (MeOH + 0.1% acetic Acid) in 5.0 min was followed by pumping 100% B for another 2 or 4 min with A being H_2_O + 0.1% acetic acid. The flow rate was 1.0 mL/min. High-resolution (positive ion) mass spectra (HRMS) were acquired using a LCMS-TOF mass spectrometer. Purity is >95% for all final compounds determined by LC-MS. Analytical HPLC was performed with prominence diode array detector (SPD-M20A). Samples were injected onto a 3.6 µm PEPTIDE XB-C18 100 Å, 150 x 4.6 mm LC column at room temperature. The flow rate was 1.0 mL/min. Various linear gradients were used with A being H_2_O + 0.1% TFA and B being acetonitrile + 0.1% TFA. Purity is >95% for key compounds in Figure 3 determined by analytical HPLC.

## (*E*)-3-(3-(3,5-Dimethylphenyl)benzo[*c*]isoxazol-5-yl)acrylic acid (1)

NaOH pellets were granulated (300 mg, 5.00 mmol) and added in 2-PrOH (7.5 mL). The mixture was put into the ultrasonic cell until NaOH was totally suspended, then was added 4-nitrobenzaldehyde glycol acetal (293 mg, 1.50 mmol). The reaction mixture was stirred until the solid was dissolve and then was added 2-(3,5-dimethylphenyl)acetonitrile (544 mg, 3.75 mmol). The reaction mixture was stirred at rt and monitored by TLC. Upon the completion of reaction, a light-yellow solid was precipitated. The solid was filtered and washed with sat. Na_2_S_2_O_3_, water and 2-PrOH.

Without further purification, the solid was added to a mixture of MeCN (6.0 mL) and water (1.5 mL), then treated with TFA (300 μL, 3.92 mmol). The reaction mixture was stirred at rt for 3 h., quenched with a saturated NaHCO_3_ solution, and extracted with CH_2_Cl_2_. The organic phase was dried (Na_2_SO_4_) and concentrated under reduced pressure. The residue was purified by ISCO silica gel column (hexane/EtOAc gradient) to afford 3-(3,5-dimethylphenyl)benzo[*c*]isoxazole-5-carbaldehyde as a yellow solid (189 mg, 0.755 mmol, 50%). ^1^H NMR (400 MHz, CDCl_3_) δ 10.01 (s, 1H), 8.38 (s, 1H), 7.83 (dd, *J* = 9.3, 1.3 Hz, 1H), 7.70-7.63 (m, 3H), 7.22 (dd, *J* = 1.6, 0.8 Hz, 1H), 2.47 (s, 6H); MS (ESI) for [M+H]^+^ (C_16_H_14_NO_2_^+^): calcd. m/z 252.10; found m/z 252.10; LC-MS: 98% purity.

To a solution of 3-(3,5-dimethylphenyl)benzo[*c*]isoxazole-5-carbaldehyde (189 mg, 0.751 mmol) in CH_2_Cl_2_ (7.5 mL) was added ethyl 2-(triphenylphosphoranylidene)acetate (274 mg, 0.788 mmol). The resulting solution was stirred at rt for 2 h. The solvent was concentrated under vacuum and the residue was purified by ISCO silica gel column (hexane/EtOAc gradient) to afford **5** as a yellow solid (237 mg, 0.738 mmol, 98%) in 98% yield and *E*/*Z* 97:3 (LCMS). MS (ESI) for [M+H]^+^ (C_20_H_20_NO_3_^+^): calcd. m/z 322.14; found m/z 322.10; LC-MS: 97% purity.

To a solution of **5** (237 mg, 0.738 mmol) in MeCN (5.0 mL) was added the aqueous solution of NaOH (88.6 mg, 2.21 mmol) in water (2.4 mL). The resulting suspension was heated at 60 °C for 1.5 h, cooled to rt, and treated with 1 M HCl to pH 1. The mixture was added small amount MeOH and refluxed for 1 h to remove the *Z*-isomer. The precipitate was filtered and washed with small amount of MeOH and water to afford the title compound as a yellow solid (183 mg, 0.625 mmol, 85%). ^1^H NMR (400 MHz, DMSO-*d*_6_) δ 8.46 (s, 1H), 7.86 (d, *J* = 9.4 Hz, 1H), 7.84-7.76 (m, 3H), 7.67 (d, *J* = 9.4 Hz, 1H), 7.24 (s, 1H), 6.61 (d, *J* = 16.0 Hz, 1H), 2.42 (s, 6H); ^13^C NMR (101 MHz, DMSO-*d*_6_) δ 167.72, 165.40, 157.19, 143.38, 139.07, 132.67, 131.48, 129.25, 127.00, 124.55, 124.20, 119.44, 115.63, 113.91, 20.84; MS (ESI) for [M+H]^+^ (C_18_H_16_NO_3_^+^): calcd. m/z 294.11; found m/z 294.10; LC-MS: 99% purity.

## (±)-*trans*-2-(3-(3,5-Dimethylphenyl)benzo[*c*]isoxazol-5-yl)cyclopropane-1-carboxylic acid (2)

To a solution of trimethylsulfoxonium iodide (139 mg, 0.630 mmol) in anhydrous DMSO (0.5 mL) was added NaH (25.2 mg, 0.630 mmol, 60% wt suspension in mineral oil) at 0 °C. After stirred 20 min at 0 °C, a solution of **5** (96.4 mg, 0.300 mmol) in anhydrous DMSO (1.0 mL) was added dropwisely. The reaction mixture was stirred at rt for 24 h and poured into a sat. NH_4_Cl solution. The mixture was extracted with EtOAc (3x), dried (Na_2_SO_4_) and concentrated under reduced pressure. The residue was purified by ISCO silica gel column (hexane/EtOAc gradient) to afford ethyl *trans*-2-(3-(3,5-dimethylphenyl)benzo[*c*]isoxazol-5-yl)cyclopropane-1-carboxylate as a yellow solid (35.3 mg, 0.105 mmol, 35%). ^1^H NMR (400 MHz, CDCl_3_) δ 7.62-7.52 (m, 4H), 7.14 (d, *J* = 2.0 Hz, 1H), 7.02 (dd, *J* = 9.3, 1.6 Hz, 1H), 4.20 (q, *J* = 7.1 Hz, 2H), 2.65-2.57 (m, 1H), 2.44 (s, 6H), 1.96 (ddd, *J* = 8.5, 5.3, 4.2 Hz, 1H), 1.64 (dt, *J* = 9.1, 5.0 Hz, 1H), 1.43-1.36 (m, 1H), 1.30 (t, *J* = 7.1 Hz, 3H); MS (ESI) for [M+H]^+^ (C_21_H_22_NO_3_^+^): calcd. m/z 336.16; found m/z 336.20; LC-MS: 95% purity.

To a solution of ethyl *trans*-2-(3-(3,5-dimethylphenyl)benzo[*c*]isoxazol-5-yl)cyclopropane-1-carboxylate (35.3 mg, 0.105 mmol) in MeCN (0.75 mL) was added a solution of NaOH (12.6 mg, 0.12 mmol) in water (0.25 mL). The resulting suspension was heated at 60 °C for 3.0 h, cooled to rt, treated with 1 M HCl to pH 1, and extracted with EtOAc (3x). The combined organic phase was washed with brine, dried (Na_2_SO_4_), and concentrated under reduced pressure. The residue was purified by reverse phase ISCO silica gel column (MeCN/H_2_O gradient) to afford **2** as a pale brown solid (21.6 mg, 70.3 μmol, 67%). ^1^H NMR (400 MHz, CDCl_3_) δ 7.63-7.52 (m, 4H), 7.14 (dt, *J* = 2.0, 1.0 Hz, 1H), 7.04 (dd, *J* = 9.1, 1.6 Hz, 1H), 2.75-2.66 (m, 1H), 2.45 (s, 6H), 1.98 (ddd, *J* = 8.4, 5.4, 4.2 Hz, 1H), 1.72 (dt, *J* = 9.6, 5.0 Hz, 1H), 1.49 (ddd, *J* = 8.4, 6.7, 4.8 Hz, 1H). ^13^C NMR (101 MHz, CDCl_3_) δ 179.00, 164.60, 157.51, 139.13, 135.52, 132.28, 130.26, 128.32, 124.52, 117.70, 116.19, 114.35, 27.47, 23.14, 21.58, 16.87; MS (ESI) for [M+H]^+^ (C_19_H_18_NO_3_^+^): calcd. m/z 308.13; found m/z 308.10; LC-MS: 97% purity.

## 3-(3,5-Dimethylphenyl)-5-(1*H*-tetrazol-5-yl)benzo[*c*]isoxazole (4)

(General procedure A) NaOH pellets were granulated (80.0 mg, 2.00 mmol) and added in 2-PrOH (1.0 mL). The mixture was put into the ultrasonic cell until NaOH was totally suspended, then was added 5-(4-nitrophenyl)-1*H*-tetrazole (38.2 mg, 0.200 mmol). The reaction mixture was stirred until the solid was dissolved, then 3,5-dimethylphenylacetonitrile (87.1 mg, 0.600 mmol) was added. The reaction was stirred at rt and monitored by TLC. Upon the completion of reaction, the suspension was acidified by glacial acetic acid to pH 5. A light-yellow precipitate was collected by filtration, washed by small amount of EtOAc, and purified by reversed ISCO chromatography to afford **4** as a yellow solid (15.1 mg, 51.8 μmol, 26%). ^1^H NMR (400 MHz, DMSO-*d*_6_) δ 8.72 (t, *J* = 1.3 Hz, 1H), 8.05 (dd, *J* = 9.3, 1.4 Hz, 1H), 7.89 (dd, *J* = 9.3, 1.0 Hz, 1H), 7.76 (d, *J* = 1.6 Hz, 2H), 7.29 (d, *J* = 1.8 Hz, 1H), 2.44 (s, 6H). ^13^C NMR (101 MHz, DMSO-*d*_6_) δ 166.39, 156.93, 154.82, 139.12, 132.94, 129.59, 126.82, 124.32, 121.15, 120.77, 116.47, 113.40, 20.89. MS (ESI) for [M+H]^+^ (C_16_H_14_N_5_O^+^): calcd. m/z 292.12; found m/z 292.10; LC-MS: 97% purity.

## 3-Phenyl-5-(1*H*-tetrazol-5-yl)benzo[*c*]isoxazole (6)

The title compound **6** (38.7 mg, 0.147 mmol, 74%) was prepared according to general procedure A from 5-(4-nitrophenyl)-1*H*-tetrazole (38.2 mg, 0.200 mmol) and benzyl cyanide (57.7 µL, 0.500 mmol) as a yellow solid. ^1^H NMR (400 MHz, DMSO-*d*_6_) δ 8.76 (t, *J* = 1.1 Hz, 1H), 8.20-8.12 (m, 2H), 8.06 (dd, *J* = 9.3, 1.5 Hz, 1H), 7.90 (dd, *J* = 9.3, 1.0 Hz, 1H), 7.77-7.61 (m, 3H); ^13^C NMR (101 MHz, DMSO-*d*_6_) δ 166.08, 156.96, 154.95, 131.43, 129.77, 129.59, 126.90, 126.76, 121.00, 120.97, 116.51, 113.56; MS (ESI) for [M+H]^+^ (C_14_H_10_N_5_O^+^): calcd. m/z 264.09; found m/z 264.10; LC-MS: 99% purity.

## 5-(1*H*-Tetrazol-5-yl)-3-(4-(trifluoromethyl)phenyl)benzo[*c*]isoxazole (7)

The title compound **7** (17.4 mg, 52.5 µmol, 26%) was prepared according to general procedure A from 5-(4-nitrophenyl)-1*H*-tetrazole (38.2 mg, 0.200 mmol) and 4-trifluoromethylphenylacetonitrile (60.2 µL, 0.400 mmol) as a yellow solid. ^1^H NMR (400 MHz, DMSO-*d*_6_) δ 8.78 (d, *J* = 1.3 Hz, 1H), 8.36 (d, *J* = 8.2 Hz, 2H), 8.11-7.99 (m, 3H), 7.96 (dd, *J* = 9.3, 1.0 Hz, 1H). ^13^C NMR (101 MHz, DMSO-*d*_6_) δ 164.30, 157.05, 130.90, 130.58, 130.38, 129.76, 127.52, 126.58 (q, *J* = 3.9 Hz), 123.80 (q, *J* = 274 Hz), 121.89, 120.54, 116.79, 114.49. MS (ESI) for [M+H]^+^ (C_15_H_9_F_3_N_5_O^+^): calcd. m/z 332.08; found m/z 332.10; LC-MS: 99% purity.

## 5-(1*H*-Tetrazol-5-yl)-3-(4-(trifluoromethoxy)phenyl)benzo[*c*]isoxazole (8)

The title compound **8** (53.5 mg, 0.154 mmol, 77%) was prepared according to general procedure A from 5-(4-nitrophenyl)-1*H*-tetrazole (38.2 mg, 0.200 mmol) and 4-trifluoromethoxyphenylacetonitrile (94.3 µL, 0.600 mmol) as a yellow solid. ^1^H NMR (400 MHz, DMSO-*d*_6_) δ 8.75 (d, *J* = 1.3 Hz, 1H), 8.34-8.25 (m, 2H), 8.08 (dd, *J* = 9.3, 1.4 Hz, 1H), 7.94 (dd, *J* = 9.4, 1.0 Hz, 1H), 7.76-7.66 (m, 2H). ^13^C NMR (101 MHz, DMSO-*d*_6_) δ 164.66, 157.01, 155.24, 150.00 (q, *J* = 1.7 Hz), 129.80, 129.07, 126.04, 122.18, 121.69, 120.56, 120.01 (q, *J* = 259 Hz), 116.55, 113.92. MS (ESI) for [M+H]^+^ (C_15_H_9_F_3_N_5_O_2_^+^): calcd. m/z 348.07; found m/z 348.05; LC-MS: 97% purity.

## 3-(4-Phenoxyphenyl)-5-(1*H*-tetrazol-5-yl)benzo[*c*]isoxazole (9)

The title compound **9** (32.0 mg, 90.0 µmol, 45%) was prepared according to general procedure A from 5-(4-nitrophenyl)-1*H*-tetrazole (38.2 mg, 0.200 mmol) and 4-phenoxyphenylacetonitrile (105 mg, 0.500 mmol) as a yellow solid. ^1^H NMR (400 MHz, DMSO-*d*_6_) 8.44 (s, 1H), 8.18 (dd, *J* = 9.3, 1.3 Hz, 1H), 8.15-8.06 (m, 2H), 7.72-7.65 (m, 1H), 7.52-7.44 (m, 2H), 7.26 (dd, *J* = 8.4, 2.1 Hz, 3H), 7.18 (dd, *J* = 7.7, 1.5 Hz, 2H); MS (ESI) for [M+H]^+^ (C_20_H_14_N_5_O_2_^+^): calcd. m/z 356.11; found m/z 356.10; LC-MS: 95% purity.

## *N*-(4-(5-(1*H*-Tetrazol-5-yl)benzo[*c*]isoxazol-3-yl)phenyl)cyclopropanecarboxamide (10)

(General procedure B) To a solution of 2-(4-aminophenyl)acetonitrile (200 mg, 1.51 mmol) in DMF (3.0 mL) was added cyclopropanecarboxylic acid (122 µL, 1.51 mmol), DIPEA (527 μL, 3.03 mmol) and HATU (633 mg, 1.66 mmol). The resulting mixture was stirred at rt for 1 h and quenched with water (2.5 mL). The product was crushed out and isolated by centrifugation. The crude product was washed with water (2x) and dried under lyophilization to afford *N*-(4- (cyanomethyl)phenyl)cyclopropanecarboxamide as an off-white solid (248 mg, 1.24 mmol, 82%). ^1^H NMR (400 MHz, CDCl_3_) δ 7.53 (d, *J* = 8.3 Hz, 2H), 7.41 (s, 1H), 7.27 (d, *J* = 8.1 Hz, 2H), 3.71 (s, 2H), 1.50 (tt, *J* = 8.0, 4.5 Hz, 1H), 1.12 – 1.06 (m, 2H), 0.87 (dt, *J* = 7.8, 3.5 Hz, 2H); MS (ESI) for [M+H]^+^ (C_12_H_13_N_2_O^+^): calcd. m/z 201.10; found m/z 201.10; LC-MS: 95% purity.

To a mixture of 5-(4-nitrophenyl)-1*H*-tetrazole (57.3 mg, 0.300 mmol) and *N*-(4- (cyanomethyl)phenyl)cyclopropanecarboxamide (50.1 mg, 0.250 mmol) in 1-butyl-3-methylimidazolium tetrafluoroborate ([Bmim]BF_4_) (2.5 mL) was added triazabicyclodecene (TBD, 348 mg, 2.50 mmol). The reaction mixture was stirred at rt and monitored by LCMS. Upon the completion of reaction, the mixture was acidified by HOAc to pH 5, quenched with brine, and extracted with EtOAc (3x). The combined organic phase was washed with brine, dried (Na_2_SO_4_) and concentrated under reduced pressure. The residue was purified by reverse phase ISCO silica gel column (MeCN/H_2_O gradient) to afford the title compound as a yellow solid (34.2 mg, 98.7 µmol, 39%). ^1^H NMR (400 MHz, DMSO-*d*_6_) δ 10.63 (s, 1H), 8.77 (s, 1H), 8.18-8.09 (m, 2H), 8.04 (dd, *J* = 9.3, 1.4 Hz, 1H), 7.96-7.89 (m, 2H), 7.87 (d, *J* = 9.4 Hz, 1H), 1.89-1.80 (m, 1H), 0.93-0.82 (m, 4H); ^13^C NMR (126 MHz, DMSO-*d*_6_) δ 172.35, 166.09, 156.94, 142.14, 133.51, 129.51, 127.66, 121.45, 121.25, 119.44, 116.34, 112.93, 40.02, 14.83, 7.68; MS (ESI) for [M+H]^+^ (C_18_H_15_N_6_O_2_^+^): calcd. m/z 347.13; found m/z 347.10; LC-MS: 98% purity.

## *N*-(3-(5-(1*H*-Tetrazol-5-yl)benzo[*c*]isoxazol-3-yl)phenyl)cyclopropanecarboxamide (11)

The title compound **11** (28.6 mg, 82.6 µmol, 33%) was prepared according to general procedure B from 2-(3-aminophenyl)acetonitrile (134 µL, 1.13 mmol) and cyclopropanecarboxylic acid (90.4 µL, 1.13 mmol) as a yellow solid. ^1^H NMR (400 MHz, DMSO-*d*_6_) δ 10.57 (s, 1H), 8.72 (s, 1H), 8.42 (s, 1H), 8.04 (d, *J* = 9.3 Hz, 1H), 7.90 (d, *J* = 9.4 Hz, 1H), 7.83 (d, *J* = 8.1 Hz, 2H), 7.62 (t, *J* = 8.0 Hz, 1H), 1.83 (t, *J* = 6.2 Hz, 1H), 0.91-0.84 (m, 4H); ^13^C NMR (214 MHz, DMSO-*d*_6_) δ 172.37, 166.10, 157.09, 155.41, 140.55, 130.41, 129.74, 127.33, 121.72, 121.33, 121.30, 120.87, 116.73, 116.68, 113.71, 14.83, 7.59; MS (ESI) for [M+H]^+^ (C_18_H_15_N_6_O_2_^+^): calcd. m/z 347.13; found m/z 347.10; LC-MS: 98% purity.

## *N*-(3-(5-(1*H*-Tetrazol-5-yl)benzo[*c*]isoxazol-3-yl)phenyl)cyclopentanecarboxamide (12)

The title compound **12** (65.4 mg, 0.155 mmol, 47%) was prepared according to general procedure B from 2-(3-aminophenyl)acetonitrile (134 µL, 1.13 mmol) and cyclopentanecarboxylic acid (122 µL, 1.13 mmol) as a yellow solid. ^1^H NMR (400 MHz, DMSO-*d*_6_) δ 10.25 (s, 1H), 8.73 (s, 1H), 8.45 (s, 1H), 8.06 (d, *J* = 9.3 Hz, 1H), 7.92 (d, *J* = 9.4 Hz, 1H), 7.85 (t, *J* = 7.8 Hz, 2H), 7.63 (t, *J* = 8.0 Hz, 1H), 2.83 (p, *J* = 7.9 Hz, 1H), 1.95-1.82 (m, 2H), 1.82-1.63 (m, 4H), 1.64-1.50 (m, 2H); ^13^C NMR (214 MHz, DMSO-*d*_6_) δ 175.14, 166.10, 163.72, 157.13, 140.72, 130.40, 129.85, 127.33, 121.83, 121.32, 120.76, 116.83, 116.68, 114.41, 113.74, 45.53, 30.19, 25.80; MS (ESI) for [M+H]^+^ (C_20_H_19_N_6_O_2_^+^): calcd. m/z 375.16; found m/z 375.10; LC-MS: 98% purity.

## 3-(3-(Piperidin-4-yloxy)phenyl)-5-(1*H*-tetrazol-5-yl)benzo[*c*]isoxazole (13)

To a solution of 2- (3-hydroxyphenyl)acetonitrile (2.00 g, 15.0 mmol), *tert*-butyl 4-hydroxypiperidine-1-carboxylate (3.33 g, 16.5 mmol) and PPh_3_ (4.33 g, 16.5 mmol) in anhydrous THF (15 mL) was added diisopropyl azodicarboxylate (DIAD, 3.28 mL, 16.5 mmol) at 0 °C. The resulting mixture was stirred at rt for 24 h. Upon completion, the reaction passed through a short pad of silica gel. The silica gel was then washed with a mixture of hexane and EtOAc (2:1, 150 mL). The organic phase was concentrated to about 50 mL and washed with a 1.0 M NaOH solution (3x) and brine. The organic layer was dried (Na_2_SO_4_) and concentrated under reduced pressure. The residue was dissolved in CH_2_Cl_2_ and loaded to silica gel column. The product was isolated from silica column chromatography as an oil, which was poured into hexane (45 mL). The mixture was sonicated until a white solid crushed out to afford a fine suspension. The precipitate was filtered and washed by hexane to afford *tert*-butyl 4-(3-(cyanomethyl)phenoxy)piperidine-1-carboxylate as a white solid (3.82 g, 12.1 mmol, 80%). ^1^H NMR (400 MHz, Chloroform-*d*) δ 7.31-7.26 (m, 1H), 6.94-6.82 (m, 3H), 4.49 (tt, *J* = 7.1, 3.5 Hz, 1H), 3.75-3.65 (m, 2H), 3.72 (s, 2H), 3.35 (ddd, *J* = 13.5, 7.6, 3.8 Hz, 2H), 1.98-1.87 (m, 2H), 1.75 (dtd, *J* = 13.8, 7.3, 3.7 Hz, 2H), 1.47 (s, 9H); MS (ESI) for [M+Na]^+^ (_18_H_24_N_2_O_3_Na^+^): calcd. m/z 339.17; found m/z 339.20; LC-MS: 96% purity.

NaOH pellets were granulated (0.98 g, 24 mmol) and added to 2-PrOH (25 mL). The mixture was stirred for 15 min before the addition of 5-(4-nitrophenyl)-1*H*-tetrazole (703 mg, 3.68 mmol). After the tetrazole was dissolved, *tert*-butyl 4-(3-(cyanomethyl)phenoxy)piperidine-1-carboxylate (775 mg, 2.45 mmol) was added. The reaction mixture was stirred at rt and monitored by LCMS. Upon the completion of reaction, to the suspension was added HOAc to pH 5. The mixture was poured into brine and extracted by EtOAc (3x). The combined organic phase was dried (Na_2_SO_4_) and concentrated under reduced pressure. The residue was purified by reverse phase ISCO silica gel column (MeCN/H_2_O gradient) to afford *tert*-butyl 4-(3-(5-(1*H*-tetrazol-5-yl)benzo[*c*]isoxazol-3-yl)phenoxy)piperidine-1-carboxylate as a yellow solid (310 mg, 0.671 mmol, 27%). ^1^H NMR (400 MHz, CD_3_OD) δ 8.72 (d, *J* = 1.6 Hz, 1H), 8.09 (d, *J* = 9.3 Hz, 1H), 7.80-7.72 (m, 2H), 7.67 (q, *J* = 1.9 Hz, 1H), 7.58 (t, *J* = 8.0 Hz, 1H), 7.23 (dd, *J* = 8.3, 2.5 Hz, 1H), 3.78 (ddd, *J* = 12.0, 7.1, 3.9 Hz, 2H), 3.41 (d, *J* = 10.7 Hz, 2H), 2.03 (dd, *J* = 13.0, 7.6 Hz, 2H), 1.76 (dtd, *J* = 12.0, 7.8, 3.8 Hz, 2H), 1.48 (s, 9H) (one proton is overlapping with CD_3_OD peak); MS (ESI) for [M+H]^+^ (C_24_H_27_N_6_O_4_^+^): calcd. m/z 463.21; found m/z 463.15; LC-MS: 98% purity.

To a solution of *tert*-butyl 4-(3-(5-(1*H*-tetrazol-5-yl)benzo[*c*]isoxazol-3-yl)phenoxy)piperidine-1-carboxylate (310 mg, 0.671 mmol) in MeOH (3.4 mL) was added a 4 N HCl solution in 1,4-dioxane (1.68 mL, 6.71 mmol). The reaction mixture was stirred at rt for 3 h and concentrated under reduced pressure. The residue was sonicated in small amount of a mixture of MeOH/H_2_O (1:1) and collected by centrifuge to afford the title compound (229 mg, 0.574 mmol, 86%). ^1^H NMR (500 MHz, CD_3_OD) δ 8.68 (s, 1H), 8.03-7.96 (m, 1H), 7.77-7.70 (m, 2H), 7.67 (d, *J* = 2.2 Hz, 1H), 7.58 (t, *J* = 8.0 Hz, 1H), 7.25 (dd, *J* = 8.2, 2.5 Hz, 1H), 3.49 (ddd, *J* = 12.8, 9.1, 3.6 Hz, 2H), 3.35-3.27 (m, 2H), 2.27 (ddd, *J* = 12.9, 9.0, 3.7 Hz, 2H), 2.13 (dq, *J* = 11.0, 3.4 Hz, 2H) (one proton is overlapping with CD_3_OD peak); ^13^C NMR (126 MHz, CD_3_OD) δ 166.21, 157.48, 157.27, 130.84, 129.12, 128.71, 121.34, 120.76, 119.74, 118.42, 116.17, 114.01, 113.85, 68.71, 40.52, 26.98 (one *sp*^*2*^ carbon is missing); MS (ESI) for [M+H]^+^ (C_19_H_19_N_6_O_2_^+^): calcd. m/z 363.16; found m/z 363.15; LC-MS: 99% purity.

## 3-(3-((3,5-Dichloropyridin-2-yl)oxy)phenyl)-5-(1*H*-tetrazol-5-yl)benzo[*c*]isoxazole (14)

To a solution of 2-(4-hydroxyphenyl)acetonitrile (66.6 mg, 0.500 mmol) in DMF (1.5 mL) was added 3,5-dichloro-2-fluoropyridine (87.1 mg, 0.525 mmol), and Cs_2_CO_3_ (326 mg, 1.00 mmol). The reaction mixture was stirred under microwave irradiation (100 W) at 120 °C for 15 min, quenched with brine, and extracted with EtOAc (3x). The combined organic phase was dried (Na_2_SO_4_) and concentrated under reduced pressure. The residue was purified by ISCO silica gel column (0-20% EtOAc/hexane gradient) to afford 2-(4-((3,5-dichloropyridin-2-yl)oxy)phenyl)acetonitrile as an off-white solid (114 mg, 0.408 mmol, 82%). ^1^H NMR (400 MHz, CDCl_3_) δ 7.96 (d, *J* = 2.4 Hz, 1H), 7.78 (d, *J* = 2.4 Hz, 1H), 7.39 (d, *J* = 8.6 Hz, 2H), 7.17 (d, *J* = 8.6 Hz, 2H), 3.78 (s, 2H); MS (ESI) for [M+H]^+^ (C_13_H_9_Cl_2_N_2_O^+^): calcd. m/z 279.01; found m/z 279.05; LC-MS: 99% purity.

The title compound **14** (19.3 mg, 45.4 µmol, 18%) was prepared according to general procedure A from 5-(4-nitrophenyl)-1*H*-tetrazole (57.3 mg, 0.300 mmol) and 2-(4-((3,5-dichloropyridin-2-yl)oxy)phenyl)acetonitrile (69.8 mg, 0.250 mmol) as a yellow solid. ^1^H NMR (400 MHz, DMSO-*d*_6_) δ 8.77 (s, 1H), 8.39 (d, *J* = 2.3 Hz, 1H), 8.21 (d, *J* = 2.3 Hz, 1H), 8.07 (dd, *J* = 9.3, 1.5 Hz, 2H), 8.00 (t, *J* = 2.0 Hz, 1H), 7.93 (d, *J* = 9.4 Hz, 1H), 7.77 (t, *J* = 8.0 Hz, 1H), 7.49 (dd, *J* = 8.1, 2.4 Hz, 1H); ^13^C NMR (101 MHz, DMSO-*d*_6_) δ 165.13, 157.01, 156.92, 154.89, 153.90, 143.77, 139.49, 131.38, 129.73, 128.27, 125.63, 124.40, 123.77, 121.24, 120.94, 119.57, 118.79, 116.60, 113.87; MS (ESI) for [M+H]^+^ (C_19_H_11_Cl_2_N_6_O_2_^+^): calcd. m/z 425.03; found m/z 425.00; LC-MS: 96% purity.

## 3-(3-(1-Cyclobutylpiperidin-4-yl)phenyl)-5-(1*H*-tetrazol-5-yl)benzo[*c*]isoxazole (15)

To a solution of 2-(3-bromophenyl)acetonitrile (600 mg, 3.06 mmol) and *tert*-butyl 4-(4,4,5,5-tetramethyl-1,3,2-dioxaborolan-2-yl)-3,6-dihydropyridine-1(2*H*)-carboxylate (994 mg, 3.21 mmol) in 1,4-dioxane (4.2 mL) and water (1.4 mL) was added K_2_CO_3_ (846 mg, 6.12 mmol). The mixture was degassed and refilled by nitrogen for three cycles. Then Pd(PPh_3_)_4_ (35.4 mg, 30.6 µmol) was added. The reaction mixture was heated under nitrogen at 90 °C for 18 h, quenched with brine, and extracted with EtOAc (3x). The combined organic phase was washed with brine, dried (Na_2_SO_4_) and concentrated under reduced pressure.

To the crude product in MeOH (10 mL) was added Pd/C (49 mg, 0.46 mmol). The reaction mixture was stirred under hydrogen atmosphere at 40 °C for 12 h, and then filtered through celite. The solvent was removed under reduce pressure, and the residue was purified by ISCO silica gel column (MeCN/H_2_O gradient) to afford *tert*-butyl 4-(3-(cyanomethyl)phenyl)piperidine-1-carboxylate as a yellow foam (751 mg, 2.50 mmol, 82%). ^1^H NMR (400 MHz, CDCl_3_) δ 7.35 – 7.28 (m, 1H), 7.20 – 7.12 (m, 3H), 4.24 (s, 2H), 3.73 (s, 2H), 2.79 (t, *J* = 12.4 Hz, 2H), 2.65 (tt, *J* = 12.1, 3.5 Hz, 1H), 1.81 (d, *J* = 13.3 Hz, 2H), 1.68 – 1.53 (m, 2H), 1.48 (s, 9H); MS (ESI) for [M+H]^+^ (C_18_H_25_N_2_O_2_^+^): calcd. m/z 301.40; found m/z 301.40; LC-MS: 98% purity.

NaOH pellets were granulated (672 mg, 16.64 mmol) and added to 2-PrOH (16 mL). The mixture was stirred for 15 min before the addition of 5-(4-nitrophenyl)-1*H*-tetrazole (382 mg, 1.99 mmol). After the tetrazole was dissolved, *tert*-butyl 4-(3-(cyanomethyl)phenyl)piperidine-1-carboxylate (500 mg, 1.66 mmol) was added. The reaction mixture was stirred at rt and monitored by TLC. Upon the completion of reaction, the suspension was diluted with MeOH and acidified with HOAc to pH 5. The reaction mixture was diluted with EtOAc. The organic phase was washed with water and brine, dried (Na_2_SO_4_), and concentrated. The residue was purified by reverse phase ISCO silica gel column (MeCN/H_2_O gradient) to afford *tert*-butyl 4-(3- (5-(1*H*-tetrazol-5-yl)benzo[*c*]isoxazol-3-yl)phenoxy)piperidine-1-carboxylate as a yellow solid (456 mg, 1.02 mmol, 61%). ^1^H NMR (400 MHz, CD_3_OD) δ 8.67 (s, 1H), 7.99 (dd, *J* = 9.4, 1.2 Hz, 1H), 7.96 – 7.89 (m, 2H), 7.72 (d, *J* = 9.4 Hz, 1H), 7.56 (t, *J* = 7.7 Hz, 1H), 7.46 (d, *J* = 7.8 Hz, 1H), 4.25 (d, *J* = 13.3 Hz, 2H), 3.30 (dt, *J* = 3.2, 1.6 Hz, 1H), 2.96 – 2.84 (m, 2H), 1.92 (d, *J* = 12.3 Hz, 2H), 1.75 – 1.63 (m, 2H), 1.48 (s, *J* = 8.1 Hz, 9H); MS (ESI) for [M+H]^+^ (C_24_H_27_N_6_O_3_^+^): calcd. m/z 447.51; found m/z 447.50; LC-MS: 95% purity.

To a solution of *tert*-butyl 4-(3-(5-(1*H*-tetrazol-5-yl)benzo[*c*]isoxazol-3-yl)phenoxy)piperidine-1-carboxylate (310 mg, 0.671 mmol) in MeOH (3.4 mL) was added a 4 N HCl solution in 1,4-dioxane (1.68 mL, 6.71 mmol). The reaction mixture was stirred at rt for 3 h and concentrated under reduced pressure. The residue was sonicated in small amount of a mixture of MeOH/H_2_O (1:1) and collected by centrifuge to afford 3-(3-(piperidin-4-yl)phenyl)-5- (1*H*-tetrazol-5-yl)benzo[c]isoxazole hydrochloride as a yellow solid (229 mg, 0.574 mmol, 86%).^1^H NMR (400 MHz, DMSO-*d*_6_) δ 8.44 (s, 1H), 8.18 (d, *J* = 8.9 Hz, 1H), 7.96 (d, *J* = 8.2 Hz, 1H), 7.90 (s, 1H), 7.73 – 7.62 (m, 2H), 7.46 (d, *J* = 7.8 Hz, 1H), 4.14 – 4.01 (m, 1H), 3.17 – 2.98 (m, 4H), 2.06 (d, *J* = 13.5 Hz, 2H), 1.96 – 1.80 (m, 2H); MS (ESI) for [M+H]^+^ (C_19_H_19_N_6_O^+^): calcd. m/z 347.40; found m/z 347.40; LC-MS: 96% purity.

To a solution of 3-(3-(piperidin-4-yl)phenyl)-5-(1*H*-tetrazol-5-yl)benzo[c]isoxazole hydrochloride (30.0 mg, 78.4 µmol), Et_3_N (21.9 µL, 0.157 mmol), and HOAc (6.7 µL, 0.12 mmol) in DMF (0.72 mL) was added cyclobutanone (17.6 µL, 0.240 mmol). The resulting mixture was stirred at rt before the addition of sodium triacetoxyborohydride (33.2 mg, 0.157 mmol). The reaction mixture was then stirred at rt for 18 h. Upon the completion, the reaction mixture was subjected to reverse phase ISCO silica gel column (MeCN/H_2_O gradient) to afford **16** as a yellow solid (31.6 mg, 67.4 µmol, 92%). ^1^H NMR (400 MHz, DMSO-*d*_*6*_) δ 8.86 (s, 1H), 8.14 – 8.00 (m, 3H), 7.91 (dd, *J* = 9.4, 0.7 Hz, 1H), 7.68 (t, *J* = 7.8 Hz, 1H), 7.52 (d, *J* = 7.8 Hz, 1H), 3.65 (dd, *J* = 16.5, 8.4 Hz, 1H), 3.46 (d, *J* = 12.1 Hz, 2H), 3.12 – 3.01 (m, 2H), 2.93 – 2.84 (m, 2H), 2.35 – 2.17 (m, 4H), 2.16 – 2.00 (m, 3H), 1.83 – 1.67 (m, 2H); ^13^C NMR (101 MHz, DMSO-*d*_6_) δ 178.55, 157.47, 146.41, 145.16, 130.62, 130.16, 129.89, 127.65, 125.68, 125.22, 121.57, 121.41, 117.03, 113.99, 58.63, 49.29, 38.91, 29.85, 25.55, 13.59; MS (ESI) for [M+H]^+^ (C_23_H_25_N_6_O^+^): calcd. m/z 401.49; found m/z 401.50; LC-MS: 97% purity.

